# Antibody escape drives emergence of diverse spike haplotypes resembling variants of concern in persistent SARS-CoV-2 infections

**DOI:** 10.1101/2025.04.17.648944

**Authors:** Luke B. Snell, Suzanne Pickering, Adela Alcolea-Medina, Helena Winstone, Jeff Seow, Carl Graham, Lorcan O’Connell, Rahul Batra, Michael H. Malim, Katie J. Doores, Gaia D. Nebbia, Jonathan Edgeworth, Stuart J.D. Neil, Rui P. Galão

**Affiliations:** Department of Infectious Diseases, School of Immunology and Microbial Sciences, King’s College London, London, United Kingdom; Centre for Clinical Infection and Diagnostics Research, Dept. of Infectious Diseases, Guy’s & St Thomas’ NHS Foundation Trust, London, UK

## Abstract

Evolution of SARS-CoV-2 in long-term persistent infections is hypothesised to be a major source of variants of concern (VOC). However, determination of viral subpopulations by linkage of intra-host variants into haplotypes has been limited by commonly used genomic sequencing techniques. We developed a sequencing and analysis methods for identifying full-length intra-host spike haplotypes and analysed their diversification over the course of persistent infections in individuals with inherited or acquired immunodeficiencies. Mutations frequently emerged in positions found in VOCs that confer escape from neutralising antibodies, while selection analyses showed strong evidence of positive selection and identified specific amino acid positions undergoing selection. In a single chronic infection lasting more than 500 days from the first wave of the pandemic, we detailed the evolution of spike as it gradually acquired mechanisms to evade both autologous and heterologous neutralising antibodies, redolent of Omicron variants. This provides one of the strongest pieces of evidence for persistent infections being the source of immune-evasive variants with accelerated evolution, underscoring their impact on the evolutionary trajectory of SARS-CoV-2.

## Introduction

The emergence and evolution of SARS-CoV-2 variants of concern (VOCs) have been key drivers of the pandemic, with successive waves of infection frequently linked to highly divergent lineages, often described as ‘saltatory’^1^. These lineages, responsible for rapid selective sweeps^2^ typified by the Alpha and Omicron waves^3^, evolve mutations that enhance receptor binding^4,5^, immune evasion^6-10^, or replication efficiency^11,12^. As a result, they exhibit increased transmissibility^13,14^ and altered disease severity^15,16^ posing significant challenges to public health measures and vaccine effectiveness^17^.

Three main theories have been proposed to explain the sudden emergence of highly divergent VOCs. The first theory suggests that mutations gradually accumulate during chains of acute infections. However, intermediate sequences leading to VOCs like Alpha or Omicron are rarely identified in global sequencing data^18^. The second hypothesis involves cross-species “spillback” transmission events, where the virus adapts to a different host before re-entering humans, as observed in mink^19^. The third and increasingly supported theory points to long-term persistent (LTP) infections in immunocompromised individuals as a key factor in VOC evolution (reviewed by Markov et al. and Sigal et al.^20,21^). These prolonged infections may create a unique environment for viral evolution, enabling sustained replication, immune-driven selection pressures and adaptations^1,22,23^. LTP infections have been observed in individuals with either inherited or acquired immunodeficiencies, with the latter arising from disease-related immunosuppression or iatrogenic causes, such as medical therapy^24-26^. Such cases often involve haematological malignancies or treatment with CD20-depleting therapies^26^.

Acute infections present strong purifying selections and narrow transmission bottlenecks that typically limit viral diversity^27,28^, while chronic infections have been suggested to enable ongoing viral replication and the emergence of highly mutated escape variants^29^. Additionally, therapeutic interventions such as monoclonal antibody and antiviral treatments may further drive viral adaptation within these immunocompromised hosts^22,30^. This is mostly evident in the viral spike protein (S), which frequently exhibits multiple amino-acid changes in domains responsible for host cell recognition and infection^24,31^, while being the primary site of diversity generation during LTP infections^21,32^. Most of these mutations cluster in key antigenic regions of spike such as the N-terminal domain (NTD) and the receptor-binding domains (RBD), and are linked to increased resistance to immune defences, particularly evasion from neutralizing antibodies^33^.

While persistent infections are recognized as potential reservoirs for VOC emergence, key questions remain about the mechanisms underlying intra-host viral evolution. The absence of transmission bottlenecks in immunocompromised individuals allows for greater intra-host diversity^34^. However, not all persistent infections result in significant genetic divergence, as some cases show little to no accumulation of mutations^35^. This highlights the need for advanced genomic approaches to better understand the natural history of persistent infections, the occurrence of adaptive mutations, the selective pressures driving these changes, and whether intra-host variation during particularly lengthy persistent infections contributes to the emergence of VOCs.

Previous genome sequencing strategies for studying the dynamic nature of SARS-CoV-2 intra-host variation during persistent infections have faced limitations^36,37^. Short-read technologies fail to link mutations into haplotypes^34^, while long-read sequencing approaches have historically suffered from high error rates, making them inaccurate for resolving intra-host variants^36,37^. To overcome these challenges, we developed a long-read high-accuracy genome sequencing workflow - achieving 99.9% of base-calling accuracy - specifically optimized for haplotyping intra-host spike populations by capturing the entire spike gene in a single read. In this study, we applied this workflow to individuals with persistent SARS-CoV-2 infections to investigate intra-host spike variation and evolution as a potential driver of VOC emergence. We then examined the interaction between antibody response and intra-host viral evolution, providing insights into the evolutionary pathways that may contribute to the emergence of future SARS-CoV-2 variants.

## METHODS

### Patient identification, sampling and definitions of persistent infection

Patients were identified at a referral centre for COVID-19 cases^26^. Cases of acute and persistent infection were identified in line with previous definitions^38^ - acute cases involved individuals without immunocompromise who were asymptomatic at the time of their initial positive test, while chronic infections were defined by PCR positivity lasting at least 30 days. Residual longitudinal nasal and throat swab samples and autologous serums from St. Thomas’ Hospital were retrieved from April 2020 to January 2024, and processed under existing ethics, which did not require patient consent and allowed linked data to be retrieved from routinely collected notes (20/SC/0310, South Central - Hampshire B Research Ethics Committee). Severity of illness was categorised as per World Health Organisation definitions^39^.

### Amplification of the spike gene sequence and high quality long-read sequencing

Nucleic acid extraction utilised the QIAGEN QIAsymphony SP system in combination with the QIAsymphony DSP Virus/Pathogen Mini Kit (Qiagen), following the off-board lysis protocol. To create amplicons spanning the full length of the spike gene sequence, nucleic acid extracts were subjected to reverse transcription polymerase chain reaction (RT-PCR) using the SuperScript™ IV One-Step RT-PCR System (Invitrogen) with primers flanking the spike region (Forward: AGGGGTACTGCTGTTATGTCTTT; Reverse: AGGCTTGTATCGGTATCGTTGC)^40^ under the following incubation conditions: 10 mins at 55.0°C; 2 mins at 98.0°C; 40 cycles of 10s at 98.0°C, 10s at 64.6°C, 2 mins 20 s at 72.0°C; followed by final extension of 5 mins at 72.0°C. PCR products were quantified using the high sensitivity dsDNA assay kit (Thermo Fisher) on the Qubit 3.0 Fluorometer (Thermo Fisher). 400 fmol of PCR products were prepared for sequencing using SQK-NBD114.24 (Oxford Nanopore Technologies, UK) according to manufacturer’s conditions with the following modifications. For the end preparation reaction 7µL of Ultra II End-prep Reaction Buffer and 3µL of Ultra II End-prep Enzyme Mix were used and incubated for 15 mins at 20°C and 15 mins at 65°C. The final library was eluted in a 30 µL volume. 20 fmol of the final library was sequenced using R10.4.1 flow cells on the GridION (Oxford Nanopore Technologies) for 24 hours.

### Bioinformatic analysis for the identification of spike haplotypes

Demultiplexing of sequencing data was performed using onboard MinKnow v23.04.5 (Oxford Nanopore Technologies). Duplex basecalling was performed using dorado v0.3.1 (https://github.com/nanoporetech/dorado) with further processing into fastq using samtools v1.10 (https://github.com/samtools/).

To identify unique spike haplotpyes, fastq files were then processed using HaploVar v1.0 (https://github.com/GSTT-CIDR, **Supplemental Methods Figure 1**). Briefly, reads with a minimum quality of Q30 (99.9% basecalling accuracy) were first filtered with Nanofilt v2.8.0 (https://github.com/wdecoster/nanofilt) and reads are identified that fully span the spike region using samtools and bedtools v.2.30.0 (https://github.com/arq5x/bedtools2), taking reads crossing both the start (21,563) and end position (25,384) of spike against Wuhan reference genome (NCBI accession MN908947.3). The identified Q30 full spike spanning reads are then subsampled from the total duplex read pool using seqtk v1.3 (https://github.com/lh3/seqtk). Bam files were created from this high quality subsampled full-length spike reads using samtools.

**Figure 1.**
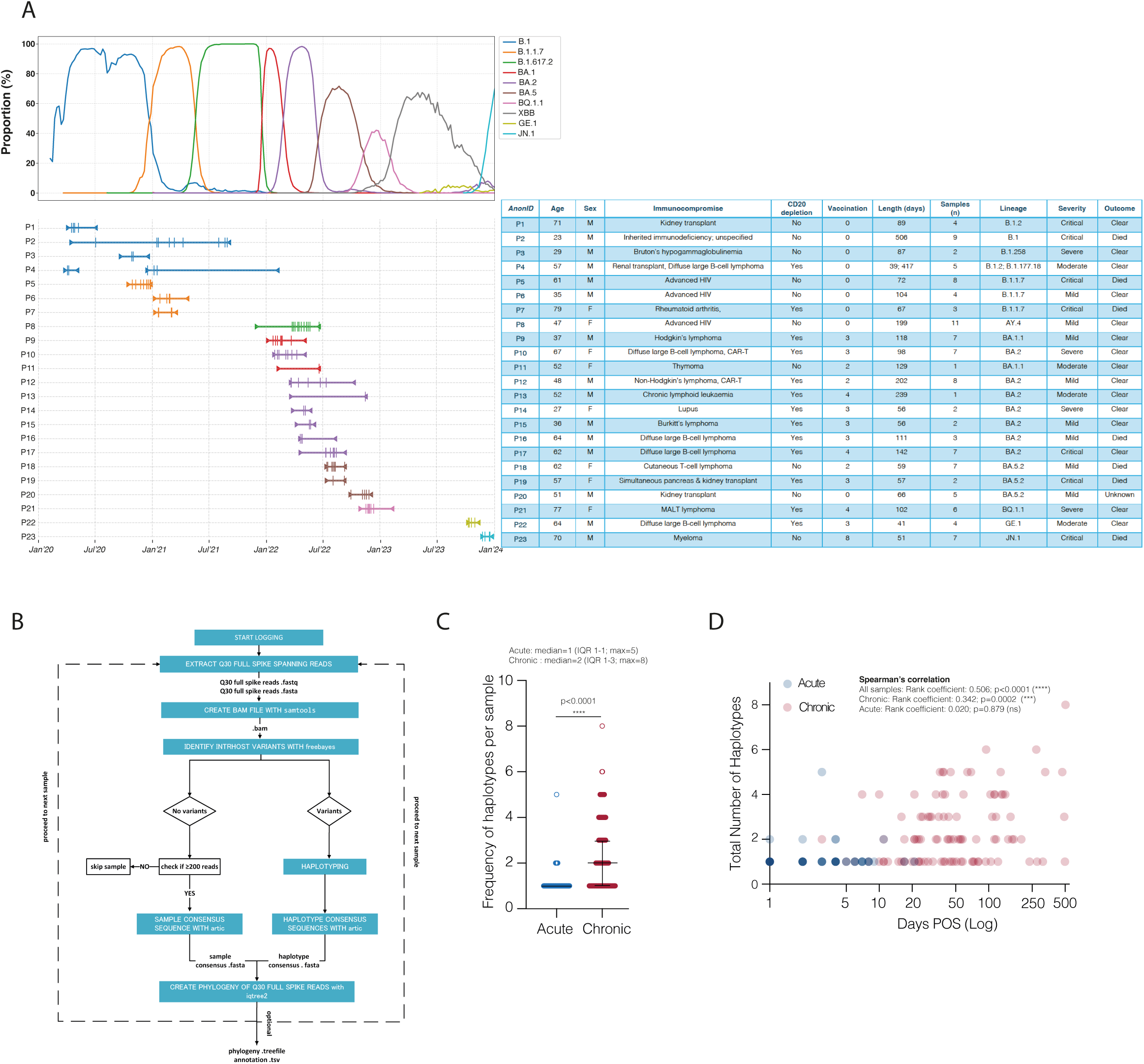
Newly developed high-accuracy long-read sequencing workflow shows diversity of SARS-CoV-2 spike haplotypes in chronic infections. **(A)** Top panel: The prevalence of major SARS-CoV-2 lineages and their descendants in England, UK, from March 2020 to January 2024, based on GISAID data. Bottom panel: Case timelines for patients with persistent SARS-CoV-2 infection, where symptom onset dates and final positive sample dates are marked with inverted triangles, and successfully sequenced samples are represented by vertical lines. Cases are colour matched to the identity of the original infecting viral lineage. Table: Overview of cases with persistent infection included in this study (see also Supplemental Table 1). **(B)** Simplified diagram showing schematic of HaploVar v1.0 bioinformatic workflow (see also Supplemental Methods Figure 1) **(C)** Number of haplotypes with a frequency greater than 5% in each sample from acutely infected individuals (blue) and chronically infected patients (red). Error bars represent mean with interquartile range across biological samples, and dots represent individual values for each sample. ****p<0.0001 as determined by Mann-Whitney test. **(D)** Scatter plot showing days post onset of symptoms (POS, Log) against the number of spike haplotypes with a frequency greater than 5% in each sample. Data points are coloured to represent acute (blue) and chronic (red) infections. Correlation determined by Spearman’s; ****p<0.0001 and ***p<0.001.

Variants (SNPs, deletions and insertions) were identified in these bam files using freebayes v1.3.7 (https://github.com/freebayes/freebayes), identifying variants at a minimum of 1% frequency in the read pool and with a minimum depth of 200 based on previous reports for identifying intra-host variants^34^. The output from freebayes was queried using awk (https://github.com/onetrueawk/awk) to identify positions of variants not fixed in the read pool to identify intra-host variants. Next, the reads were further processed to identify haplotypes, specifically how the identified intra-host variants exist in linkage with each other in each read. To achieve this, the Q30 full-length spike reads were converted to fasta format using seqtk (https://github.com/lh3/seqtk) before being mapped against the Wuhan reference genome using gofasta v1.2.1 (https://github.com/virus-evolution/gofasta). Reads with ambiguous mapping were discarded. For each full-length spike read, the nucleotide at each position of intra-host variation was extracted, discarding the rest of the read where the positions were invariant, giving a haplotype for each read. Unique haplotypes within the read pool were then identified, the frequency of the occurrence of each haplotype in the read pool was counted, and reads were separated into discrete read pools according to haplotype using seqtk. Consensus sequences for each haplotype were then created using a modified version of the ARTIC bioinformatic pipeline reflecting the custom primer scheme and a minimum coverage depth of 10x.

### Testing ability of the spike haplotyping workflow to determine minority variants

To allow determination of viral copy number a standard curve was established using a quantified N-gene from a Wuhan SARS-CoV-2 genome (TIB Molbiol 30-7454-71) and applying a RT-qPCR method (NEB Luna® SARS-CoV-2 RT-qPCR Multiplex Assay Kit). Absolute viral copy number was determined for B.1.1.7 (National Institute for Biological Standards and Control, #101019) and B.1.351 lineage (#101022) before artificial mixing 10^4^ genome copies with minority variant at 0%, 0.5%, 1%, 2%, 5%, 10%, 20%, before testing with the spike haplotyping workflow.

### Phylogenetic and selection analysis

Maximum likelihood phylogenies were derived using IQ-TREE v2.3.0^41^ fitted using a GTR+G4+I model, and with branch testing utilising Shimodaira–Hasegawa-like approximate likelihood-ratio test with 1000 replicates. Trees were further processed in ete3 v3.1.3 (https://anaconda.org/conda-forge/ete3) for display. Molecular-clock phylogenies were constructed in TimeTree v0.11.3 (https://github.com/neherlab/treetime).

Rates of synonymous and non-synonymous mutation were calculated using the Nei-Gojobori method (Jukes Cantor model) in Mega11^42^ assuming the standard genetic code. Variation in rates were assumed to be gamma distributed (shape parameter = 1.0) across the spike gene region and gaps were treated as pairwise deletions. Ambiguous positions were not included in comparisons. For each patient, the reference haplotype used for comparison in these calculations was the most abundant haplotype in the first successfully sequenced sample.

Tests were performed to identify which sites were under selective pressure using HYPHY v2.5.62^43^. Mixed Effects Model of Evolution (MEME)^44^ was used to test sites under episodic selection pressure both overall and at specific timepoints. Fixed effects likelihood (Contrast-FEL)^45^ was used to determine which sites have significantly different rates of change between timepoints. Phylogenic resampling was set at 1000 replicates. Molecular clocks utilized TempEST v1.5.3 (https://tree.bio.ed.ac.uk/software/tempest/).

When evaluating mutations in haplotypes, the lineages considered for comparison were any assigned a Greek letter by the World Health Organisation (i.e Alpha, Beta, Gamma, Delta, Episilon, Kappa, Eta, Iota, Lambda, Epsilon, Theta, Zeta, Mu, Omicron) and predominant lineages that came after Greek letters were assigned (BA.1-5, JN.1 and KP.3).

### Cells

HEK293T/17 (ATCC CRL 11268TM) and Vero-E6 (ATCC CRL 1586TM) cells were obtained from the American Type Culture Collection. Vero-E6 cells were modified to stably express TMPRSS2 via lentiviral vector transduction^10^. HeLa-ACE2 cells were generously provided by James E. Voss. All cell lines were cultured in DMEM supplemented with GlutaMAX (Gibco, UK) and 10% FCS, maintained at 37°C with 5% CO^2^.

### Pseudovirus production

Sub-confluent HEK293T/17 cells were transfected in 10 cm dishes with 2 µg of the HIV-1 8.91 Gag-Pol plasmid, 3 µg of the CSXW (HIV-firefly luciferase) plasmid, and 2 µg of the SARS-CoV-2 spike plasmid using 35 µg of PEI-Max (1 mg/ml, Polysciences). Cells were incubated at 37°C, and the medium was replaced 6–12 hours post-transfection. Pseudoviruses were harvested 72 hours after transfection and filtered through 0.45 µm filters. Low-titre preparations were concentrated via ultracentrifugation through a sucrose cushion. To achieve this, pseudoviruses were initially treated with 10 U/mL of recombinant DNase I (Merck) in the presence of 10 µM MgCl₂ for 2 hours at 37°C. The DNase-treated preparations were then layered over a 20% sucrose cushion in PBS and ultracentrifuged at 28,000 rpm for 1 hour and 30 minutes. After centrifugation, the supernatant was removed, and the pellets were resuspended in serum-free DMEM GlutaMAX.

All spike plasmids used in this study were codon-optimized and included full-length cytoplasmic tails. The SARS-CoV-2 B.1 spike plasmid was generously provided by Prof. Nigel Temperton. Longitudinal spikes representing the most common haplotypes from patient 2 were synthesized by GenScript.

### Viruses

The UK SARS-CoV-2 Wave 1 (lineage B.1) reference strain, England 02 (England 02/2020/407073), was obtained from Public Health England. Full-length viruses were isolated from patient nasal and throat swab samples by diluting 200 µl of the sample in 1.5 ml of DMEM GlutaMAX containing 2% FCS, filtering through 0.45 µm filters, and infecting Vero-E6-TMPRSS2 cells. Supernatants were collected upon the appearance of visible cytopathic effects (CPE). For virus propagation, 100 µl of the isolated virus was added to confluent Vero-E6-TMPRSS2 cells cultured in 75 cm² flasks with DMEM GlutaMAX supplemented with 2% FCS. Cells were monitored daily, and cultures were harvested upon visible CPE. The harvested cultures were filtered through 0.45 µm filters, aliquoted, and stored at -80°C until use.

### Plaque Assays

To determine viral titres, plaque assays were conducted in 6-well plates. Virus samples were serially diluted 10-fold, and 500 µl of each dilution was added per well to confluent Vero-E6-TMPRSS2 cells, followed by incubation at 37°C for 1 hour. After incubation, 500 µl of a pre-warmed overlay (0.1% agarose in DMEM GlutaMAX supplemented with 2% FCS) was added to each well, and the plates were incubated at 37°C for 72 hours. Cells were fixed with 4% formaldehyde at room temperature for 30 minutes, then stained with 0.05% crystal violet in ethanol for 5 minutes. Wells were washed with PBS, air-dried, and plaques were counted. Viral titres were calculated by averaging the results from three independent assays.

### Neutralisation assays

Neutralisation assays were performed using longitudinal serum samples from patient 2, as well as wave 1 representative sera^46^, and monoclonal antibodies (mAbs) representative of different competing binding groups^47,48^, and the commercial mAbs sotrovimab, imdevimab and casirivimab.

Neutralization assays with pseudovirus were conducted using HeLa-ACE2 cells^48^. mAbs or serum (heat-inactivated at 56°C for 30 minutes before initial use) were serially diluted in DMEM GlutaMAX (Gibco, UK) and incubated with pseudovirus at 37°C for 1 hour. HeLa-ACE2 cells were prepared at a concentration of 5 × 10⁵ cells/mL, and 50 µl (2.5 × 10⁴ cells/well) was added to each well. Plates were incubated for 72 hours at 37°C before measuring luciferase activity using the Steady-Glo® Luciferase Assay System (Promega, UK) with a VICTOR™ X Multilabel Reader (Perkin Elmer). For neutralisation assays with full-length SARS-CoV-2 infectious virus, mini plaque reduction neutralisation tests (PRNT) were performed^48^. Vero-E6-TMPRSS2 cells were seeded the day before infection at a density of 30,000 cells per well of a 96-well plate in DMEM supplemented with 2% FCS. Sera were serially diluted in DMEM GlutaMAX (Gibco, UK) and incubated with virus (optimised to achieve 80-200 plaques per well) at 37°C for 1 hour (50ul of virus was added to 50ul of diluted sera) before adding to cells. This was incubated for a further 1 hour at 37°C before the addition of pre-warmed carboxymethylcellulose overlay (Sigma-Aldrich, C4888) to a final concentration of 0.5%. Plates were incubated 37°C for 16-20 hours before removing supernatant and fixing with 4% formaldehyde in PBS (30 minutes at room temperature). Fixed cells were then washed in PBS, permeabilised in 0.2% Triton X-100 for 15 minutes at room temperature, then blocked with 3% milk in PBS for 15 minutes at room temperature before addition of murinised anti-nucleocapsid antibody CR3009 (2ug/ml) for 45 minutes at room temperature. Plates were washed twice with PBS and incubated for a further 30 minutes at room temperature with secondary antibody goat anti-mouse IgG (Fc-specific)-peroxidase (Sigma A2554; 2 ug/ml). Plates were washed twice with PBS before addition of TrueBlue peroxidase substrate (50ul per well; SeraCare 50-78-02). Plates were incubated for 20-60 minutes until clear, dark blue plaques were visible, before removal of substrate and air drying. Plaques were identified and counted using an AID EliSpot Reader with EliSpot 8.0 software.

### Statistics

Statistical analyses were performed using both GraphPad Prism v10, and SciPy v1.9.3 in python v3.1.1 (https://pypi.org/project/scipy/), with a significance level set at p < 0.05.

## RESULTS

### High-accuracy long-read sequencing workflow defines diversity of SARS-CoV-2 full-spike haplotypes in long-term persistent infections

To investigate the dynamic nature of SARS-CoV-2 intra-host variation during such prolonged infections, we identified 23 chronically infected patients (defined as having PCR positive tests for a minimum of 30 days) with diverse underlying immunocompromised states such as immunosuppressive therapies, advanced HIV, autoimmune disorders or cancer-related immunodeficiencies. These patients exhibited varied histories of vaccination and/or SARS-CoV-2 treatments (**Figure 1A**, **Supplemental Table 1**). For this study, we collected 123 convenience longitudinal nasal and throat swabs samples between April 2020 and January 2024 from these immunocompromised patients. Patients were persistently infected with various SARS-CoV-2 variants for durations ranging from 39 to 506 days (median=93.5 days; IQR 58-139 days), with a median of 5 samples per patient successfully sequenced (IQR 2-7 samples) (**Figure 1A**). 69 samples collected during the same period from individuals with acute SARS-CoV-2 infections were used as control group (**Supplemental Material Database**). The lineage distribution among the acute samples included 1 A, 16 B.1, 11 Alpha, 3 Delta, 3 BA.2, 1 BA.4, 14 BA.5 and 20 XBB lineages.

To resolve low-frequency intra-host single nucleotide variants (iSNVs), we developed a high-accuracy nanopore-based long-read workflow capable of spanning the entire SARS-CoV-2 spike coding region. This workflow produces Q30 reads using R10.4.1 flow cells (Oxford Nanopore Technologies), giving a 99.9% base-calling accuracy comparable to Illumina short-read technology, while simultaneously enabling haplotyping through its long-read capabilities. The bioinformatic workflow “HaploVar 1.0” was designed to extract Q30 reads spanning the entirety of spike gene sequence, before identification of intra-host variants (see Methods) by haplotyping and using the ARTIC pipeline to generate consensus sequences of identified haplotypes. Maximum likelihood and molecular clock phylogenies were constructed from the haplotype consensus sequences (**Figure 1B, Supplemental Methods Figure 1**).

To test the performance of the haplotyping workflow in identifying and quantifying intra-host variants, triplicate tests using artificial mixtures containing 10^4^ copies of viral RNA from Alpha and Beta lineages demonstrated that the workflow consistently detected single integer minority populations (**Supplementary Methods Figure 2A**). Furthermore, haplotypes above 5% were reproducibly identified with as few as 200 Q30 full-spike reads upon down-sampling of sequencing data from clinical specimens (**Supplemental Methods Table 1)**, in accordance with others that also use a depth of 200 reads for calling intra-host variants^34^. The lower limit of detection of the workflow for achieving the threshold of ≥200 Q30 reads required for successful haplotyping was 100 viral genome RNA copies (**Supplementary Methods Figure 2B**). Replicate sequencing of chronic clinical samples showed that haplotypes comprising more than 5% of the total were reproducibly detected (**Supplemental Methods Figures 2C, Supplemental Methods Table 2**). Therefore, a 5% minimum frequency threshold was applied for reporting of haplotypes throughout the study, which is similar to the 3% thresholds applied in Illumina sequencing^35^.

**Figure 2.**
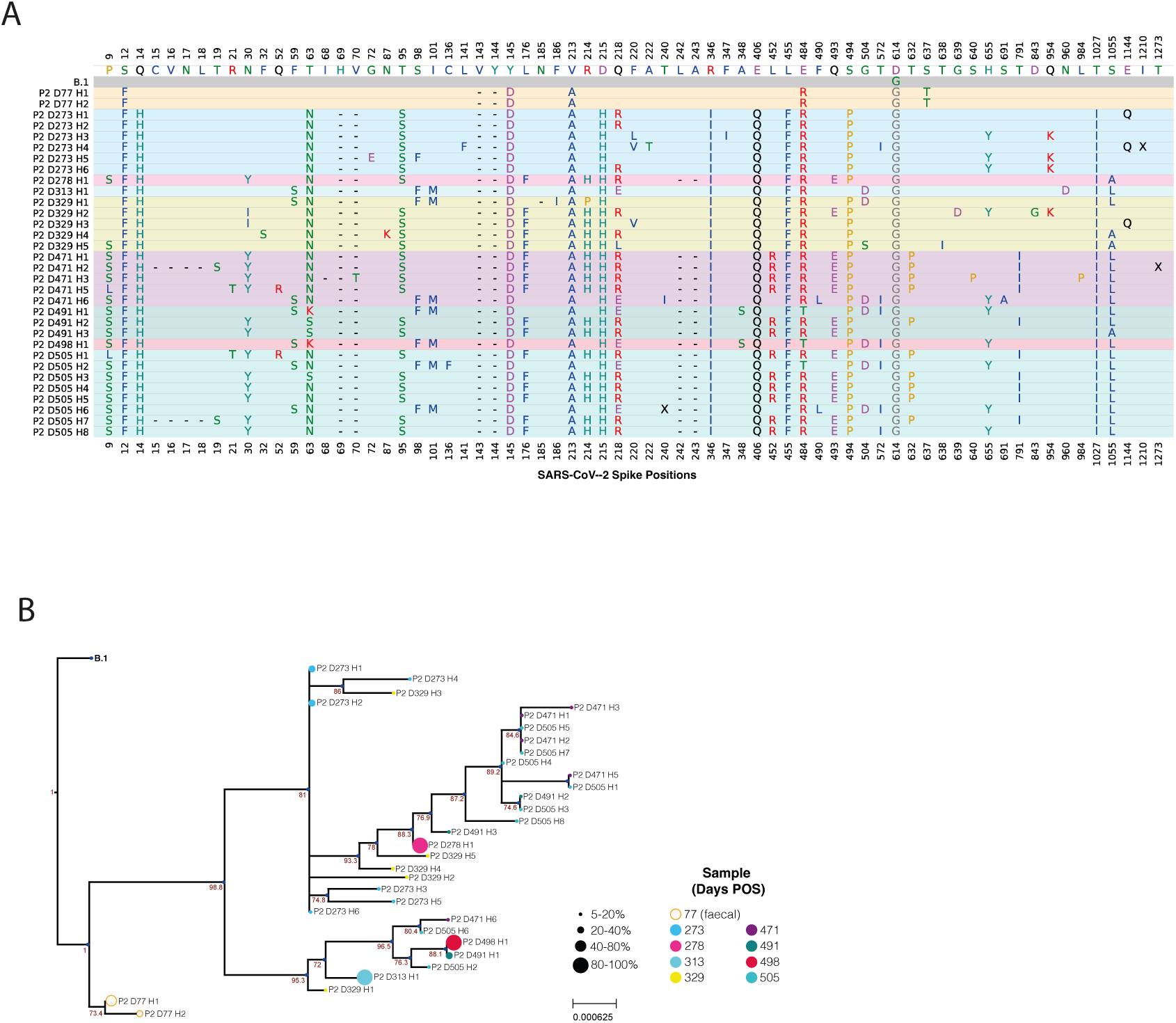
Example of intra-host evolution of full-length spikes during SARS-CoV-2 chronic infection. **(A)** Non-synonymous mutations identified in haplotypes from longitudinal samples collected from Patient 2. Samples are displayed chronologically from the earliest to the latest, with each sample background uniquely shaded. Haplotype notation included the patient identifier (P), day of sampling post-onset of infection (D), and the haplotype rank in the sample (H). Haplotypes are ordered from highest to lowest proportion. For positions with non-synonymous mutations, the amino acid residue from the ancestral SARS-CoV-2 reference sequence (NCBI MN908947.3) is listed first, followed by the corresponding residue(s) in the infecting variant. Non-synonymous mutations present in the infecting variant that remain unchanged during chronic infection are highlighted in grey. Positions not called are represented by an “X”. **(B)** A divergence phylogeny of each haplotype representing at least 5% of total haplotypes identified at each longitudinally collected sample from a single persistent infection (P2), with the infecting variant designated as the outgroup. Tip labels are colour-coded based on their corresponding sample timepoints, and their sizes are scaled to reflect the relative proportion of the haplotype within each sample. Branches are labelled with SH-like approximate likelihood ratio test results, with branches below 70% support collapsed.

**Table 1.** List of VOC-defining and VOC-like mutations observed in spike haplotypes from patients with long-term persistent infections. See also supplemental material database.

| Anon. ID | Haplotypes (number) | VOC-like mutations | VOC lineage-defining mutations |
| --- | --- | --- | --- |
| P1 | 7 |  |  |
| P2 | 32 | L8H, T19S, T95S, V213A, D215H, R346I, E484R, E484T, F490L, Q493E, Q954K | Q52R (Eta), 69/70- (Alpha), 143/144- (BA.1), 242/243- (Beta), L455F (XBB), L452R (Delta), H655Y (Omicron), T1027I (Gamma) |
| P3 | 8 | A243V | 69/70- (Alpha), T1027I (Gamma) |
| P4 | 9 |  | E484K (Beta) |
| P5 | 19 | Y144V, W152L |  |
| P6 | 8 |  | A67V (Eta), T95I (BA.1), D215G (Mu), E484K (Beta) |
| P7 | 5 |  | S371F (BA.2), N440K (BA.2), E484Q (Kappa), A701V (Beta) |
| P8 | 30 | L8K, S477I, D796H | K417T (Gamma), N440K (BA.2), S477N (BA.1), Q493E (KP.3), P621S (JN.1) |
| P9 | 18 | S371I, N764E | S371L (BA.1), S365F (BA.1), K417N (Beta), K417T (Gamma), N460K (JN.1) |
| P10 | 10 | K417I | K356T (JN.1) |
| P11 | 5 |  | S371L (BA.1), S365F (BA.1), K417N (Beta) |
| P12 | 12 | K356R |  |
| P13 | 1 |  |  |
| P14 | 3 | D796H |  |
| P15 | 2 | Y248H |  |
| P16 | 6 |  |  |
| P17 | 20 | R346I, K356R, E484V |  |
| P18 | 14 |  |  |
| P19 | 2 | A570T | 144- (BA.1) |
| P20 | 19 | E156K, S371-, S373-, S375A, S375S, T376P/T | 144- (BA.1) |
| P21 | 11 |  | 69/70- (Alpha) |
| P22 | 9 |  |  |
| P23 | 14 |  |  |

**Table 2.**
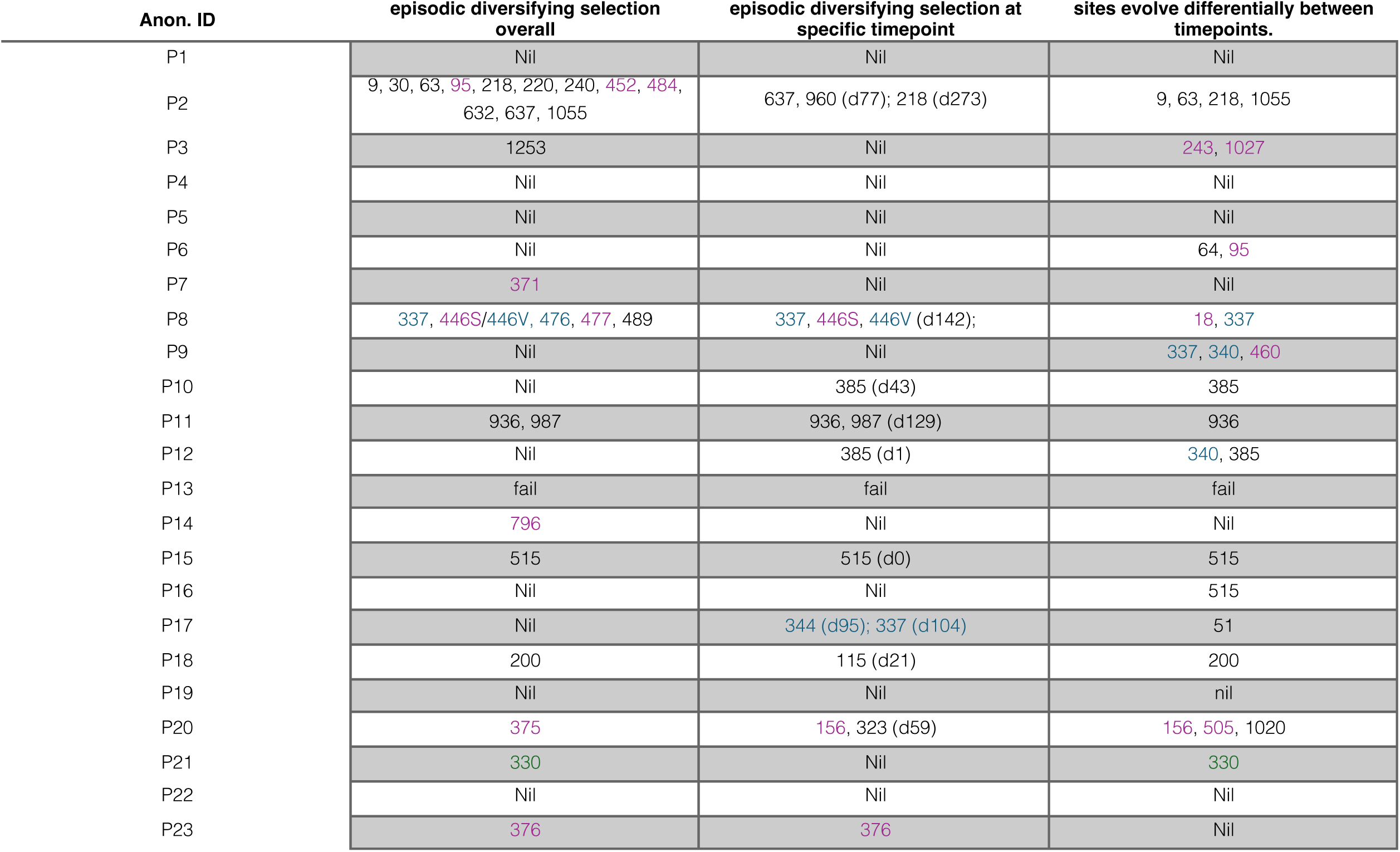
Results of phylogenetic testing to identify amino acid positions under episodic diversifying selection overall, at specific timepoints, and sites evolving differentially over time. Positions are coloured if VOC-associated (purple), reported as conferring resistance to monoclonal antibodies (blue), or previously described to be associated with persistent infections (green).

This high-accuracy workflow was then employed to explore how SARS-CoV-2 intra-host variation evolved during the course of acute and chronic infections in our panel of infected individuals (**Figure 1A**, **Supplemental Material Database**). Of the 219 samples analysed using the spike haplotype workflow, 69 of 96 acute samples and 115 of 123 chronic samples met quality control (QC) criteria. The Ct values for samples that passed and failed QC were both normally distributed (Shapiro-Wilk test: p = 0.227 and p = 0.907, respectively), with samples passing QC having significantly lower Ct values compared to those that failed (median = 18.0, SD = 4.1 vs median = 23.0, SD = 4.4; t-test: p < 0.0001) (**Supplemental Figure 1A**). For acute samples, the median number of haplotypes exceeding 5% frequency per sample was 1 (IQR: 1-1, max: 5), while chronic samples had a median of 2 haplotypes (IQR: 1-4, max: 8) among samples collected at least 30 days post-symptom onset, indicative of chronic infection (**Figure 1C**).

Correlations were assessed between the number of haplotypes and number of Q30 full spike reads or Ct values to ensure that haplotype counts were not an artefact of sequencing workflow. As expected, there was no significant correlation between the number of Q30 full spike reads and haplotypes (Spearman’s rank coefficient (r) = -0.119; p = 0.107) (**Supplemental Figure 1B**), ruling out sequencing depth as a confounding factor. Similarly, no correlation was observed between the number of haplotypes and the Ct value of the sample (r = 0.114; p = 0.134) (**Supplemental Figure 1C**). Most importantly, the number of haplotypes in each sample showed a significative correlation with days post onset of symptoms (r = 0.506; p<0.0001), which was naturally most prominent in samples from chronic infections (r = 0.342; p = 0.0002) when compared with acute samples (r = 0.020; p = 0.879) (**Figure 1D**). As final step in the workflow, spike haplotypes with a minimum 5% representation were used to construct both divergence and molecular clock phylogenies. A representative analysis for Patient 2 is illustrated in **Figure 2A-B**, while similar analyses for all patient samples are provided in **Supplemental Figure 2A-W**. In summary, we developed a new high-accuracy long-read workflow able to resolve intra-host spike haplotypes present at low frequency, while demonstrating not only that diversity correlates with the length of infection, but also that this is a highly dynamic process.

### Analysis of spike haplotypes from chronic infections reveals accelerated evolution rates, signatures of positive selection, and changes in residues associated with VOCs

Haplotypes present in samples were analysed for non-synonymous (ns) mutations in the spike gene, relative to the SARS-CoV-2 reference sequence and the infecting lineage. A moderate positive correlation was observed between the number of ns spike mutations and days post-infection (**Figure 3A**, r = 0.62; p < 0.0001). Furthermore, the rate of spike-specific mutations in our cohort of persistently infected individuals exceeded that observed in the global population, with molecular clock analysis revealing cases such as P2, P4, P11 and P23 where evolutionary rates were considerably higher than those of contemporaneous VOCs (**Supplemental Table 2**). There was no correlation between the rate of spike evolution and the timing of chronic infections during the pandemic (r= -0.062, p=0.784) **(Figure 3B**). This suggests that the evolutionary patterns of chronic spikes have remained relatively consistent over time and faster than that seen in acute infection.

**Figure 3.**
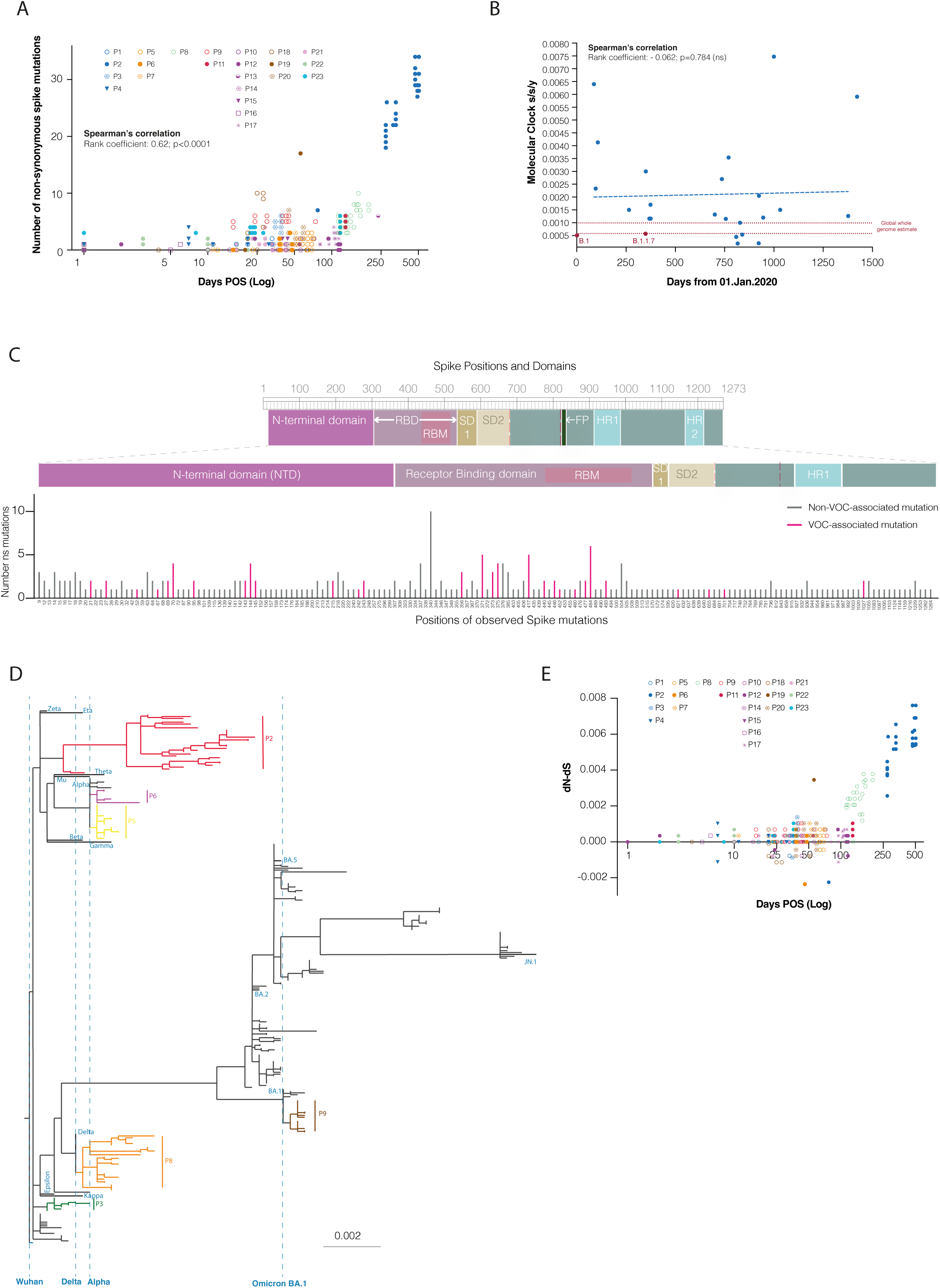
Evolution on spike haplotypes during long-term infections is characterised by changes in VOC-associated residues, accelerated evolutionary rates, and events of positive selection. **(A)** Scatter plot depicting the number of mutations found in each spike haplotype against days post-onset of symptoms (log). Individual patients have unique markers. Correlation determined by Spearman’s; ***p<0.001. **(B)** Scatter plot illustrating the correlation between the molecular clock (s/s/y) evolutionary rates estimated for the infecting variant spike of each patient (blue) or selected VOC spikes (red) against the date of infection, measured in days from January 01, 2020. Evolutionary rate intervals estimated for global whole genomes are represented by red dashes lines. Correlation determined by Spearman’s; ns>0.05. See also Supplemental Table 2. **(C)** Graphical representation of SARS-CoV-2 spike gene and corresponding protein domains within (top panel). Number of with non-synonymous mutations observed at each amino acid position in the spike protein from all patients. Patients with multiple non-synonymous mutations at the same position are counted more than once. Positions without any mutations identified are absent. Bars are coloured pink where the position corresponds to a lineage-defining mutation in VOC or VOC-associated mutation (bottom panel). **(D)** Divergence phylogeny of spike haplotypes retrieved longitudinally from persistent infections and global VOC spike. Selected patients are coloured, and certain VOC lineages are labelled. The ancestral SARS-CoV-2 reference sequence (NCBI MN908947.3) is designated as the outgroup. Branches with SH-like approximate likelihood ratio test results below 70% are collapsed. **(E)** Scatter plot depicting the difference of non-synonymous substitutions to synonymous substitutions (dN - dS) calculated for each haplotype retrieved from each persistent infection compared to the most abundant haplotype in the patients first successfully sequenced sample, plotted against the number of days post onset of symptoms (Log). Individual patients have unique markers.

Analysis of all spike haplotype populations identified in our panel of chronic patients revealed that non-synonymous mutations were primarily concentrated in highly antigenic regions such as the receptor binding domain (RBD) and, to a lesser extent, the N-terminal domain (NTD) (**Figure 3C**). Additionally, haplotypes were analysed for the presence of lineage-defining VOC mutations or mutations at spike positions that are altered in VOCs (‘VOC-associated mutations’). For this evaluation, lineages assigned a Greek letter by the World Health Organization were included for comparison, along with subsequent lineages of significant epidemiological importance (**Table 1**, **Supplemental Material Database**). While a few cases showed no VOC lineage-defining mutations (P1, P13, P16, P18, P22 and P23), the majority acquired multiple VOC-defining mutations (e.g., P2: n=7; P6, P7: n=4; P8, P9: n=5) and/or VOC-like mutations (e.g. P2: n=11; P20: n=4) (**Table 1**, **Figure 3C, Supplemental Material Database**). Notably, a comparison with global SARS-CoV-2 spike diversity revealed extensive intra-host evolution in persistent infections, with some branch lengths exceeding the divergence observed between the Wuhan reference sequence and extant VOCs (**Figure 3D**). This is particularly evident with P2, where the patient was initially infected with the B.1 lineage and, during the length of a single chronic infection, viral spikes subsequently evolved a genomic distance nearing that of an Omicron BA.1 spike and exceeding that of a BA.2 spike (**Figure 2A**, **Figure 3D**).

Haplotypes were further analysed for evidence of positive selection driving their evolution. Longitudinal analysis revealed that 65% (156/240) of haplotypes had a positive dN-dS value, indicating positive selection, while 24% had no mutations (**Figure 3E**, **Supplemental Material Database**). Phylogenetic analysis identified 36 spike positions under episodic diversifying selection across 11 patients (**Table 2**). Sixteen of these positions were associated with VOCs, suggesting frequent emergence and positive selection of VOC-like mutations in persistent infections. Notably, 1/36 position (P330S) was previously linked to chronic infections^49^, while 5/36 positions (P337, E340, A344, G446, G476) were associated with resistance to monoclonal antibodies.

Based on this observation, we analysed spike haplotypes from patients in our cohort who received monoclonal antibody treatment during their chronic infection to examine how mutations evolved in targeted regions (**Figure 4A-C**). Among the ten patients treated with sotrovimab, seven acquired mutations associated with evasion at one or more spike positions - P337, E340, R346 and/or K356 (**Figure 4A**). P17 exhibited three further mutations on the defined sotrovimab footprint that were not previously associated with resistance to this monoclonal antibody – N334K, A344S and R357K (**Figure 4A-C**). Additionally, two of three patients treated with casirivimab and imdevimab developed known resistance mutations within the corresponding binding regions (**Figure 4A-C**). P8 exemplified how resistance mutations can evolve following mAb treatment during chronic infections. This patient received casirivimab and imdevimab on day 98 post-onset of symptoms and by day 113, several resistance mutations (E406A, N440K, V445A, G446S/V, Y453F) had already emerged. However, as this is the earliest available sample for this patient, it can’t be excluded that at least some of these mutations were already present. Additional resistance mutations, such as K417T, E484Q and Q493E, appeared later in infection (**Figure 4D**). Furthermore, on day 128 the same patient was treated with sotrovimab, and known resistance mutations P337L and E340A were detected soon after on days 142 and 151, respectively. The emergence of multiple resistance mutations, none of which were detected in untreated patients, strongly suggests that monoclonal antibody therapy can quickly drive the selection of escape mutations in persistent infections.

**Figure 4.**
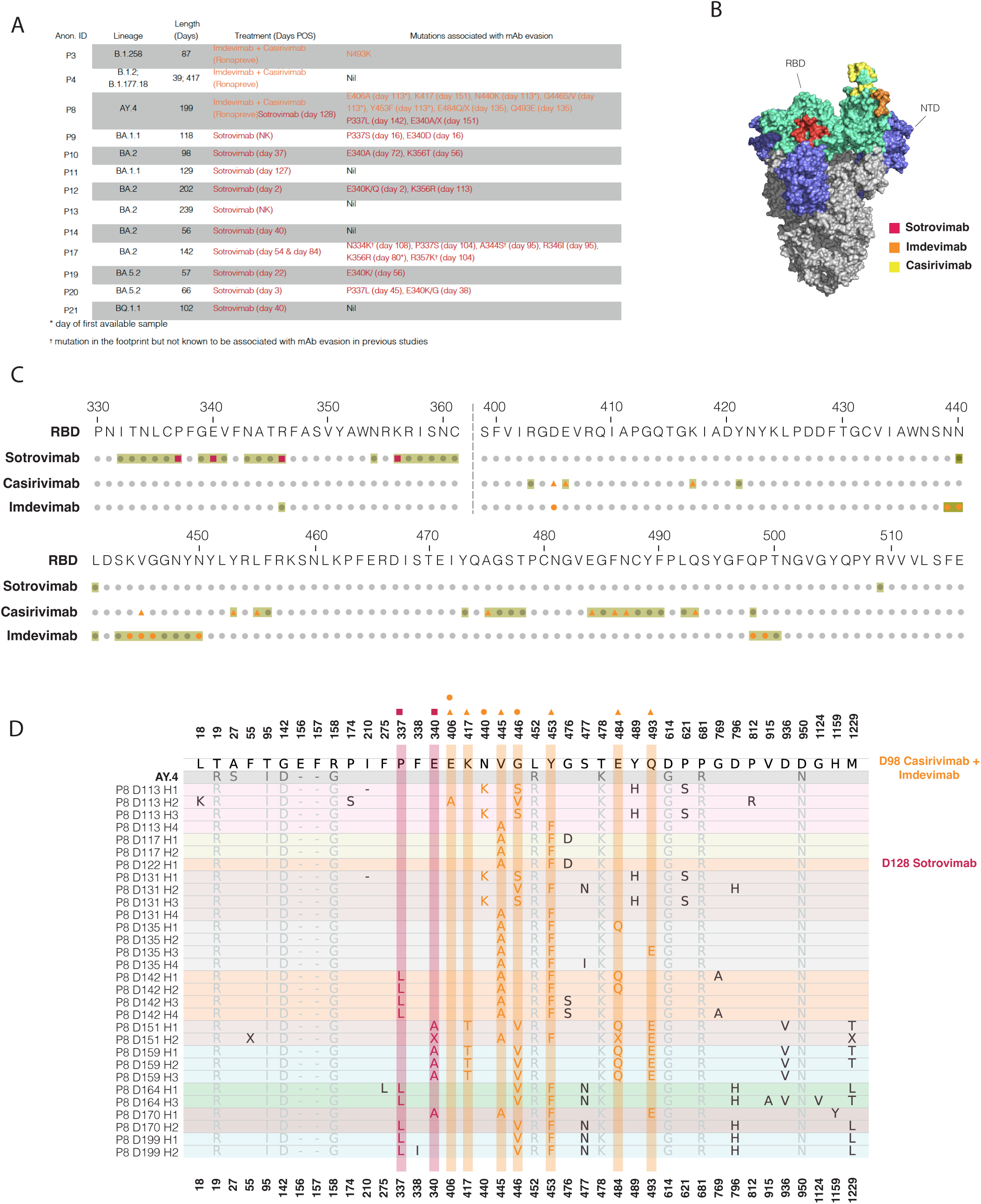
Mutations linked to monoclonal antibody (mAb) evasion can rapidly emerge following treatment in chronically infected patients. **(A)** Details of clinical cases involving long-term persistently infected patients treated with sotrovimab (red) and/or combinations of imdevimab and casirivimab (orange), along with identified mutations linked to evasion from these monoclonal antibodies (mAb). Mutations present on the day of the first available sample are indicated with (*), while those observed within the mAb binding footprint but not yet associated with mAb evasion are marked with (†). **(B)** Structure of full spike (S) of SARS-CoV-2 showing binding footprints for sotrovimab (red), casirivimab (REGN10933, yellow) together with imdevimab (REGN10987, orange). RBD and NTD domains of spike are labelled in cyan and blue, respectively. mAb footprints for sotrovimab were determined using pdb ID: 6WPS, and the Regeneron mAbs with pdb ID : 6XDG. **(C)** The full RBD sequence of ancestral SARS-CoV-2 (NCBI MN908947.3), with monoclonal antibody (mAb) binding footprints highlighted in light green. Mutations detected in spike haplotypes from patients treated with mAbs are marked in red for sotrovimab and in orange for the casirivimab and imdevimab combination. **(D)** Non-synonymous mutations identified in haplotypes from longitudinal samples collected from Patient 8. Samples are displayed chronologically from the earliest to the latest, with each sample background uniquely shaded. Dates of first treatment with mAbs are indicated on the right as days post-onset of symptoms. Haplotype notation included the patient identifier (P), day of sampling post-onset of infection (D), and the haplotype rank in the sample (H). Haplotypes are ordered from highest to lowest proportion. For positions with non-synonymous mutations, the amino acid residue from the Wuhan reference sequence (NCBI MN908947.3) is listed first, followed by the corresponding residue(s) in the infecting variant. Non-synonymous mutations on mAb footprints are highlighted in red for sotrovimab and in orange for casirivimab and imdevimab combination.

Collectively, these findings provide compelling evidence that elevated evolutionary rates occur during chronic infections, in part driven by events of positive selection, while supporting the hypothesis that long-term persistent infections play a key role in the evolution of variants of concern.

### Immune pressure leads to an evolving universal evasion from wave 1 neutralising responses

To directly assess intra-host immune pressures that may contribute to the evolution of spike, we performed detailed studies on longitudinal samples obtained from one individual with an exceptionally long disease course (P2 in **Figure 1A and Supplemental Figure 1A**). This individual became symptomatic and tested positive for SARS-CoV-2 in April 2020, admitted to hospital in June 2020 (day 77) with a persistent infection with lineage B.1 and treated with remdesivir. They were declined for compassionate treatment with antibody-based therapies. The patient passed away in September 2021, on day 505 of the infection, with multiple co-morbidities and overlapping acute illnesses (**Figure 1A**).

Spike genes were cloned from nasal swabs obtained at regular time points throughout infection (77, 118, 178, 266, 329 and 505 days post infection; **Figure 5A**). Samples were not available from this individual prior to day 77, therefore for the purposes of comparison the day 0 (infecting) virus was assumed to be an ancestral D614G sequence which represented the majority known spike sequence circulating at the time in the UK. Haplotype analysis from this individual demonstrated increasing diversification of viral sequences over time, resulting in 5 and 8 spike haplotypes above 5% representativity of the total quasispecies identified at days 329 and 505 respectively (**Figures 2A-B**). For these time points the 5 most abundant haplotypes were cloned. In parallel, full-length infectious virus was successfully cultured from a day 329 nasal swab.

**Figure 5.**
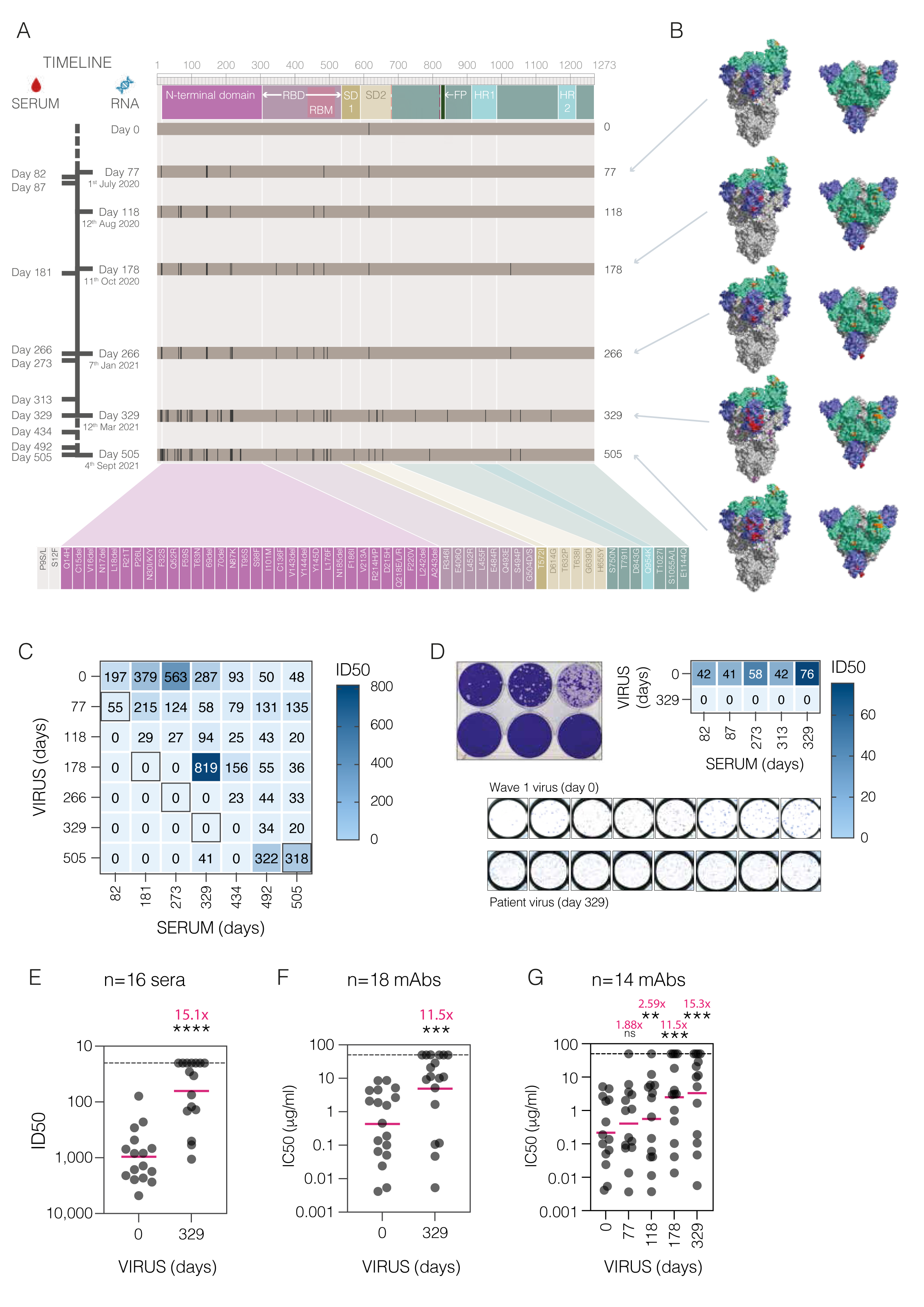
Longitudinal characterisation of intrahost SARS-CoV-2 spike evolution and humoral immune evasion during a long-term chronic infection in one individual. **(A)** Schematic representation of SARS-CoV-2 spike sequence evolution in one individual over the course of a 505-day infection. A timeline of serum samples and nasal swabs from this individual is shown on the left as days post-infection, with a schematic of the spike protein including all major domains at the top. The accumulation of individual amino acid mutations is shown over time and mapped to spike protein domains. The identity of each of these mutations is listed at the bottom, coloured according to the spike domain in which they occur. **B)** Trimeric spike structures showing the location of accumulating amino acid mutations. RBD is shown in green, NTD in blue. RBD mutations are highlighted in orange, NTD in red and visible mutations in other parts of spike are shown in purple. **(C)** Heatmap of autologous longitudinal humoral immune response over the course of a chronic SARS-CoV-2 infection in one individual. Neutralising responses were assessed using lentiviral vectors pseudotyped with sequential SARS-CoV-2 spike proteins from days 0, 77, 118, 178, 266, 329 (P2 D329 H3 In Figure 2A) and 505 (P2 D505 H1 in Figure 2A) post infection (shown in **(A)**) and longitudinal autologous serum samples from days 82, 181, 273, 329, 434, 492 and 505. Each square shows reciprocal mean neutralising titres (ID50) for a given pseudovirus-serum pair, derived from two independent experiments, with colour intensity proportional to neutralisation potency. Boxed numbers represent mean ID50 values for contemporaneous virus-serum pairs (or where not available, the closest possible match). **(D)** Infectious virus was isolated from a day 329 nasal swab, with plaque morphology shown in the left panel. Neutralisation of the day 0 ancestral virus was compared with the day 329 virus using sequential autologous serum samples from days 82, 87, 273, 313 and 329 post-infection. Each square shows reciprocal mean neutralising titres (ID50) for a given pseudovirus-serum pair, derived from two independent experiments, with colour intensity proportional to neutralisation potency. An example of results seen in the mini plaque reduction assay is shown in the bottom panel. **(E)** Neutralisation of spike from assumed infecting virus (day 0; ancestral D614G) and day 329 spike-pseudotyped virus by early acute sera (10-31 DPOS) from individuals infected in wave 1 of the pandemic. Each point represents the mean ID50 value for a given serum sample, derived from two independent experiments, with the pink line indicating the geometric mean titre (GMT) for n=16 sera. Neutralising titres of the two pseudoviruses were compared using a two-tailed paired Wilcoxon signed rank test (p<0.0001, ****). Numbers in pink indicate the fold change in GMT between the groups. **(F)** Neutralisation of day 0 and day 329 spikes was assessed using n=18 monoclonal antibodies. Each point is the mean IC50 value for a given mAb, derived from two independent experiments. The pink line shows the overall GMT for all mAbs against each spike, with numbers in pink indicating the fold change in GMT. Significance was determined by two-tailed Wilcoxon signed rank test (p=0.0001, ***). **(G)** 14 monoclonal antibodies were used for more detailed stepwise escape from wave 1 monoclonal antibodies using spikes from days 0, 77, 118, 178 and 329. Pink lines indicate GMT for n=14 mAbs against each spike, with numbers in pink showing the fold change in GMT between each spike and the ancestral (day 0) spike. Significance was determined by two-tailed Wilcoxon signed rank test (day 77 p=0.54; day 118 p=0.0085**; day 178 p=0.0004***; day 329 p=0.0006, ***).

A total of 55 amino acid mutations were detected in the spike protein throughout the course of infection in this one individual (**Figure 5A** and **Supplementary Figure 3**A). The majority of these mutations occurred in antigenic regions of spike (**Figure 5B**), with 32 in the NTD and 8 in the RBD. Several of the mutations overlapped with known VOC lineage-defining mutations or VOC-like mutations (**Table 1**), with the clustering reminiscent of that seen for Omicron BA.1 (**Supplementary Figure 3**B).

Neutralisation assays were performed with lentiviral vectors pseudotyped with spikes from sequential time points and longitudinal autologous serum samples, as close to contemporaneous as available (**Figure 5A** and **C**). Overall, titres were low compared with typical titres seen in immunocompetent cohorts (e.g. Dupont et al.^46^), yet all serum samples tested were able to neutralise the D614G ancestral variant (**Figure 5C**, Day 0 virus). A shifting window of immune escape was seen, with serum samples that preceded the spike isolation timepoint having no neutralising effect, contemporaneous serum-spike pairs showing weak or no neutralisation, and serum samples later than the spike timepoint increasing in neutralising potency, peaking at approximately 100-150 days after the time of spike isolation (**Figure 5C**). The day 505 spike was the exception to this rule, with contemporaneous sera neutralising the most common spike haplotype from this timepoint. Results with autologous sera were confirmed using full-length virus isolated from a day 329 nasal swab (**Figure 5D**). While all serum samples tested (days 82, 87, 273, 313 and 329) were able to neutralise the day 0 virus (ancestral B.1), none of them neutralised the day 329 virus. Together these results clearly illustrate a stepwise evolution of spike driven by the humoral immune response, resulting in spikes that are completely refractory to neutralisation by earlier autologous antibodies.

To assess whether the neutralisation escape characterised in later spikes was specific only to the autologous neutralising response or represented a generalised escape from typical wave 1 responses, 16 heterologous serum samples from acute wave 1 infection were compared for neutralisation potency against the ancestral D614G (day 0) and day 329 spikes (**Figure 5E**). Neutralisation titres against the day 329 spike were significantly lower than the D614G ancestral (day 0) spike, with an overall 15.1-fold difference between the geometric mean titres (GMTs). Thus, a spike protein isolated late in a long-term infection demonstrates significant immune escape from both autologous and heterologous humoral immune responses, analogous to that seen for later SARS-CoV-2 saltatory variants such as BA.1.

To probe the escape from wave 1 neutralisation in more detail, a panel of monoclonal antibodies - isolated from individuals infected during wave 1^50^, or naïve individuals vaccinated with ancestral spike-based vaccine^48^ - were tested against lentiviral vectors bearing spikes from P2. 18 mAbs were tested in total, comprising a range of potencies and binding specificities. Similar to results seen with wave 1 serum samples, neutralisation of the day 329 spike was significantly weaker than the D614G ancestral (day 0) spike, with an 11.5-fold difference in GMT (**Figure 5F**). This was further dissected to assess stepwise escape, using spikes from days 77, 118, 178 and 329 (**Figure 5G**). The accumulation of individual RBD mutations over time (**Figure 5A** and **Supplementary Figure 3**A) led to an incremental neutralisation escape, with each time point accounting for a shift in GMT (**Figure 5G**), with a bigger change occurring between day 118 and 178 due to two RBD mutations (R346I and E406Q). Separating the results into individual mAb binding groups allowed a more precise mapping of the effects of these mutations on neutralisation (**Supplementary Figure 3**C). In particular, we observed consistent evasion of Group 4 mAbs over time, which were the most common binding group isolated from vaccinated individuals^48^. Commercial mAbs, used therapeutically prior to the post-Omicron era, provided additional opportunities to map details of immune escape, given that several of these have been crystalised in complex with the SARS-CoV-2 spike. While the effects of imdevimab and sotrovimab fluctuated over time, the overall trend was for the P2 spikes to remain susceptible to neutralisation by these mAbs (**Supplementary Figure 3**D), in contrast to the effects seen with the Group 4 mAbs in **Supplementary Figure 3**C. Interestingly, casivirimab was rendered completely ineffective by day 178. The escape can be attributed to the acquisition of an E484R mutation from day 77 onwards, L455F mutation from day 118 onwards, then a E406Q mutation at day 178, all of which are located in the casirivimab footprint^51^. The commercial mAbs were also tested with multiple haplotypes from two timepoints (days 329 and 505), confirming that haplotypes from the same timepoints gave broadly similar results for the different mAbs.

In sum, we have dissected the evolution of spike in a single, exceptionally long-term SARS-CoV-2 infection and provided clear evidence that intra-host humoral immune pressure drives the characteristic accumulation of spike mutations seen in novel VOCs. We have mapped out a pathway whereby highly mutated viruses can arise in one individual, beginning with infection by an ancestral virus in the first wave of the pandemic and culminating in a variant that is broadly similar in its mutation profile and immune evasion properties to Omicron 3 months prior to the latter’s emergence in southern Africa.

## Discussion

This work describes the development of a novel long-read sequencing workflow integrated with the new bioinformatic tool, HaploVar 1.0, designed here for detecting and characterizing diverse spike haplotypes that emerge during long-term persistent infections. Haplotypes exhibited significant divergence and signs of positive selection, with many sharing key spike mutations found in variants of concern (VOCs). We then used these data to conduct phenotypic characterisation of these spike haplotypes, in what we believe to be one of the longest recorded persistent SARS-CoV-2 infection. Through these studies, we mapped out a pathway whereby a highly mutated variant can arise in a single individual, driven by escape from the autologous antibody response but resulting in a generalised resistance to first wave neutralising responses. Compared to previous studies, our approach provides a more comprehensive view of intra-host viral diversity and population dynamics over time, while also allowing for the assessment of positive selection and uniquely enabling the linkage between genotype and phenotype.

As shown in this study, persistent SARS-CoV-2 infections can lead to viral sequences that exhibit an unusually high number of mutations compared to the original infecting strain, indicating accelerated evolution relative to acute infections. These sequences show an overrepresentation of non-synonymous substitutions, sometimes with several amino acid changes occurring at the same site, as well as small deletions and insertions, suggesting rapid adaptation within chronically infected individuals rather than neutral diversification^24,29,38,52^. During acute infection transmission chains, where each infection is brief, viral populations undergo severe transmission bottlenecks^27,53^. However, intra-host transmission between cells typically lacks such constraints, allowing continuous viral replication and providing opportunities for adaptive mutations to accumulate and eventually become dominant. The rate of SARS-CoV-2 evolution in chronically infected individuals varies (reviewed in ^21^) but here we demonstrate cases can display mutation rates comparable to or exceeding those leading to saltatory emergence of VOCs, while other cases of persistent of persistent infection exhibit a slower evolutionary pace similar to contemporaneous circulating lineages. These differing findings may reflect the inherent challenges in interpreting consensus sequences from persistent infections, as such sequences might not capture the full complexity of the viral population. Depending on the prevalence of co-existing variants, consensus sequences may resemble the original strain more closely, potentially underestimating the full range of viral diversity. The sequencing workflow developed in this study addresses these limitations, offering a more precise depiction of the dynamic intra-host evolution of the spike protein during persistent infections.

Our study substantiates the theory that long-term persistent infections can drive the emergence of VOCs and underscores how studying these individuals may provide valuable insights to predict future developments in the SARS-CoV-2 pandemic that may also be of relevance to the selection of antigenic drift variants in other endemic respiratory viruses. We specifically note that, in certain cases of prolonged infection, the evolutionary trajectories of the identified spike haplotypes exhibit long branch lengths, akin to those seen in contemporaneously emerging VOCs such as Beta, Delta and Omicron. For example, spike proteins from P2 contained a total of 55 possible mutations, including seven VOC-defining mutations and 11 VOC-like mutations. Phylogenetic analysis further indicates that many of these positions are under diversifying selection.

SARS-CoV-2 is well-documented to evolve known neutralising antibody escape mutations during persistent infections^34,49,52,54^, but here our unique access to longitudinal paired viral sequences and serum samples over the course of an exceptionally long-persistent infection has allowed us to document the stepwise creation of a highly evolved Omicron-like variant. We show a that a shifting window of evasion from weak neutralising responses in an immunocompromised individual can result in a virus with universally reduced sensitivity to first wave antibody responses from immunocompetent individuals, in the form of heterologous sera, monoclonal antibodies isolated from vaccinees and infected individuals, and a commercial therapeutic monoclonal antibody. Interestingly, this is achieved with relatively few mutations in the RBD, including some that have been more recently associated with recent Omicron sub-variants. For example, L455F, known for forming the so-called FLiP mutations with F456L, which have recently appeared in XBB lineages and are recognized for their role in immune evasion and antibody escape^55^, as well as mutations at R346 and Q493E. However, in contrast to VOCs and particularly the Omicron sub-variants, in which the majority of mutations arise in the RBD, we observed a larger number of deletions and substitutions in the NTD of the P2 spikes. The potential effects of these on the conformation of the spike and exposure of key neutralising epitopes remain to be characterised. Additionally, we observed spike haplotypes in P2 carrying the H655Y mutation, a hallmark of the Gamma and Omicron VOCs, which has been linked to enhanced viral replication, changes in spike protein cleavage and altered cell entry pathways^56^, facilitating transmission^57^. Remarkably, spike haplotypes with mutations now recognized as critical in the pandemic had already evolved by the time the final sample from P2’s long-term persistent infection was collected in September 2021, three months before the emergence of the Omicron BA.1 variant. This highlights the potential of studies like this to identify and characterise key mutations that might evolve in future VOCs.

Selective pressure in long-term persistent infection can also be externally driven, for example, through the administration of convalescent plasma^54^ or monoclonal antibodies^34,58^, as demonstrated in this study for the latter. In line with earlier findings^59^, we observed swift viral adaptations following treatments with monoclonal antibodies. This was marked by the frequent emergence of well-documented escape-related mutations, including P337L and E340 – linked to resistance against sotrovimab – as well as E484Q and Q493E, associated with escape from imdevimab and casirivimab.

To definitely demonstrate the role of persistent infections in the emergence of VOCs, it is crucial to show that VOC-like variants arising in such contexts are capable of onward transmission. The transmission advantage of SARS-CoV-2 VOCs has been linked not only to adaptive immune escape, but also other immune evasion mechanisms, increased infectivity, altered cell entry processes, and changes in spike protein cleavage and structure. Although our findings show that mutations do occur on P2 spike haplotypes at sites among those inferred to have positive effects on SARS-CoV-2 VOCs transmission^60^ – such as L455F, H655Y and modifications in R346 and Q954 – there are several of the most prominent that didn’t evolve such as N460K and Q498R associated with enhanced fusogenicity and spike processing, increased ACE2 binding affinity and transmissibility^61-63^, or P681H/R suggested enhance spike cleavage and shown to confer a level of resistance to IFN-beta^10,64^. Together these might reflect that while highly divergent lineages may be transmitted^65,66^, they often fail to sustain transmission at the population level^67^. The relative rarity of these population saltation events may be linked to distinct selective pressures driving evolution, as mutations selected for during persistent infections have been proposed to favour intra-host rather than inter-host transmission^68^. Interestingly, our selection analyses has also identified previously unreported loci, offering potential insights into adaptation during host-pathogen interactions.

While different types or degrees of immunocompromise can possibly lead to divergent evolutionary patterns, we found no evidence that host factors such as immunocompromise, age, sex, vaccination status, or virus lineage influenced evolutionary rates. Overall, viral adaptation to the immune system appeared to be the primary driving force. However, a limitation of this study, together with its retrospective nature, is that only nasopharyngeal swabs were analysed, which may have led to an underestimation of viral population diversity by overlooking subpopulations present in the deeper airways or the gut.

Future studies should explore the role of specific mutation constellations in shaping viral phenotypes, including immune escape and fitness. Investigating factors such as HLA typing and T-cell responses could provide deeper insights into the mechanisms driving viral evolution, while characterizing non-spike mutations arising during chronic infections may shed light on innate immune adaptations that support viral persistence. To date, limited research has examined these dynamics, but the long-read sequencing workflow presented here offers a powerful tool for studying the phenotypic effects of linked mutations. These insights could enhance our understanding of how mutations interact to influence viral evolution and transmission.

## Resource availability

### Lead contact

Further information and requests for resources and reagents should be directed to and will be fulfilled by the lead contact, Rui Pedro Galao.

### Materials availability

All requests for resources and reagents should be directed to the lead contact. There are restrictions to the availability of some of the clinical samples used in this study due to their scarcity or the lack of remaining material. This includes nasopharyngeal swabs and sera. All further unique/stable reagents generated in this study will be made available on request after completion of a materials transfer agreement (MTA).

### Data and code availability

Sequencing data is available at the NCBI Sequencing Read Archive (BioProject PRJNA1247580), while code for HaploVar v1.0 can be found in https://github.com/GSTT-CIDR.

## Supporting information

Supplemental Material Database

## Acknowledgments

We are extremely grateful to all patients and staff at St Thomas’ Hospital who participated in this study. We thank all colleagues in the G2P consortia for the insightful discussions, as well as Prof Emma Thompson, Dr Ana Da Silva Filipe, and teams for review of initial genomic findings. With thanks to Prof Sergei Pond for his advice on performing phylogenetic selection analysis.

This research was funded by the UK Medical Research Council Discovery Awards to the G2P/G2P2 consortia (MC/PC/15068 and MR/Y004205 to MHM, KD, SJDN, RPG) and the Wellcome Trust to the G2P-global consortium (226141/Z/22/Z to MHM, KD, SJDN), and the Huo Family Foundation. This study was further supported by the UK Department of Health via an NIHR Comprehensive Biomedical Research Centre award to Guy’s and St Thomas’ NHS Foundation Trust in partnership with King’s College London and King’s College Hospital NHS Foundation Trust. SJDN was supported by a Wellcome Trust Senior Fellowship (WT098049AIA). MHM was supported by a Wellcome Trust awards (106223/Z/14/Z and 222433/Z/21/Z). LBS was supported by UK Medical Research Council Fellowship (MR/W025140/1). The funding sources of this study had no influence in study design, data collection and analysis, data interpretation, or the preparation of the report, or in the decision to submit this manuscript for publication.

## Author Contributions

Planning and conceptualisation: L.B.S., S.P., K.J.D., G.D.N., J.E., S.J.D.N., R.P.G.

Investigation, methodology and data analysis: L.B.S., S.P., A.A-M, H.W., J.S., C.G., L.O’C., R.P.G.

Funding acquisition: L.B.S., R.B., M.H.M., K.J.D., G.D.N., J.E., S.J.D.N., R.P.G.

Software: L.B.S.

Resources: K.J.D., J.E.

Writing manuscript: L.B.S., S.P., S.J.D.N., R.P.G.

Review and editing manuscript: L.B.S., S.P., A.A-M, H.W., J.S., C.G., L.O’C., R.B., M.H.M., K.J.D., G.D.N., J.E., S.J.D.N., R.P.G.

## Declarations of interest

Jonathan Edgeworth is employed part time as VP Medical Affairs by Oxford Nanopore Technologies

**Supplemental Figure 1. (A)** Histogram showing Ct value of samples successfully haplotyped (blue) compared to those that failed haplotyping (red). Normality of passed and failed samples was tested using Shapiro-Wilk; p>0.05 confirms the hypothesis of normality. **(B, C)** Scatter plots showing number of spike haplotypes with a frequency greater than 5% in each sample used in this study against logarithmic transformed number of Q30 positive full-length spike reads (B), or Ct values (C).

**Supplemental Figure 2. (A-W)** Amino-acid alignment matrices representing non-synonymous mutations identified in haplotypes from longitudinal samples collected from Patients 1-23 (top panels). Divergence phylogeny trees and molecular clock trees of haplotypes collected longitudinally from the same single persistent infections (bottom panels). See also Figure 2.

**Supplemental Figure 3. Detailed analyses of RBD mutations over time and sequential escape from mAbs**

**(A)** Detailed breakdown of the individual spike amino acid mutations shown in **Figure 5A**. Each amino acid mutation is listed. Spike haplotypes for days 329 and 505 are listed in order of frequency.

**(B)** Graphical comparison of the location of all observed mutations in P2 spike proteins with those found in the Omicron BA.1 spike. Red bars indicate Identical positions of mutations.

**(C)** Neutralisation of spike from assumed infecting virus (day 0; ancestral D614G) and days 77, 118, 178 and 329, grouped according to mAb binding specificity. Each point is the mean IC50 value for a given mAb and spike combination, derived from two independent experiments. The key shows amino acid mutations in the RBD for each longitudinal spike.

**(D)** In depth neutralisation assays of longitudinal spike proteins from days 0, 77, 117, 178, 329 and 505, including multiple spike haplotypes for the day 329 and 505 time points. Each point is mean IC50 value derived from at least two independent experiments.

**Supplemental Table 1.** Clinical details of the cases of long-term persistent infection included in this study.

**Supplemental Table 2.** Molecular clock (s/s/y) evolutionary rates estimated for the infecting variant spike of each patient or selected VOC spikes^34,69^. Where the R^2^ is greater than 0.75 the difference in rate compared to contemporaneous rate is given.

**Supplemental Methods Figure 1.** Diagram showing schematic of HaploVar v1.0 bioinformatic workflow.

**Supplemental Methods Figure 2.**. **(A)** Table and logarithmic scatter plot displaying the detection, in triplicate, of pre-defined minority populations of spike haplotypes using the HaploVar 1.0 workflow on empirical mixes containing 10^5^ copies of viral RNA from Alpha and Beta lineages. **(B)** Determination of the minimum total number of viral genome RNA copies required to obtain at least 200 Q30-positive full-length reads for the application of HaploVar 1.0. A table and scatter plot illustrate the reproducibility of detecting Q30-positive full-length reads (n=4) in samples with total viral genome RNA copies ranging from 100 to 1,000 **(C)** Logarithmic scatter plot displaying proportion of identified full-spike haplotypes on clinical samples (n=9) in two independent replicates (see also Supplemental Methods Table 2). Correlations in **A and C** determined by Pearson’s; ****p<0.0001.

**Supplemental Methods Table 1.** Impact of down-sampling on the reproducibility of spike haplotype determinations in clinical samples. HaploVar v1.0 workflow was used on five clinical samples to obtain high-quality Q30-positive full-spike reads, which were then serially diluted down to 200 Q30-positive reads. The reproducibility of determining spike haplotype frequencies was subsequently assessed for all samples and dilutions.

**Supplemental Methods Table 2.** Replicate identification and quantification of full-spike haplotypes in clinical samples (n=9)

**Supplemental Material Database. (1)** Line list of all samples processed. **(2)** List of VOC mutations. **(3)** List of mutations found in spike haplotypes for each patient. **(4)** Synonymous and non-synonymous rates in each haplotype.

## Notes

https://github.com/GSTT-CIDR

