## Supplemental Figures and Tables for "Antibody escape drives emergence of diverse spike haplotypes resembling variants of concern in persistent SARS-CoV-2 infections"

Supplemental Figure 1

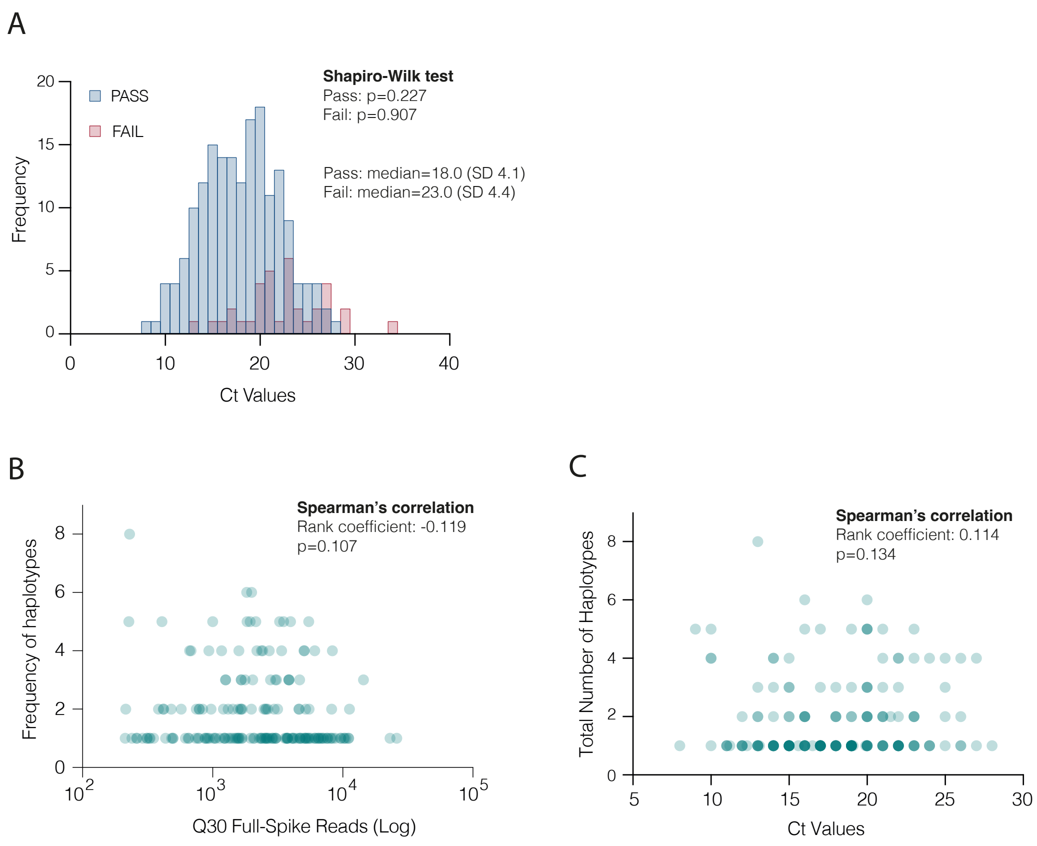

Supplemental Table 1

| Anon. ID | Age | Sex | Immunocompromise | CD20 depletion | Vaccin. (N) | Date Symptoms Onset | Date First Positive | Date Last Positive | Length (days) | Samples (N) | Lineage | Severity | Outcome | Treatment |
|---|---|---|---|---|---|---|---|---|---|---|---|---|---|---|
| P1 | 71 | M | Kidney transplant | No | 0 | 2020.04.06 | 2020.04.06 | 2020.07.03 | 89 | 4 | B.1.2 | Critical | Clear | Remdesivir (Date NA/TBC) |
| P2 | 23 | M | Inherited immunodeficiency; unspecified | No | 0 | 2020.04.17 | 2020.04.19 | 2021.09.04 | 506 | 9 | B.1 | Critical | Died | Remdesivir (Date NA/TBC) |
| P3 | 29 | M | Bruton's hypogammaglobulinemia | No | 0 | 2020.09.22 | 2020.10.05 | 2020.12.17 | 87 | 2 | B.1.258 | Severe | Clear | Ronapreve, Remdesivir (Date NA/TBC) |
| P4 | 57 | M | Renal transplant, Diffuse large B-cell lymphoma | Yes | 0 | 2020.03.28; 2020.12.16 | 2020.03.28; 2020.12.16 | 2020.05.05; 2021.02.05 | 39; 417 | 5 | B.1.2; B.1.177.18 | Moderate | Clear | Ronapreve, Remdesivir (Date NA/TBC) |
| P5 | 61 | M | Advanced HIV | No | 0 | 2021.01.03 | 2021.01.14 | 2021.03.15 | 72 | 8 | B.1.1.7 | Critical | Died | Remdesivir (Date NA/TBC) |
| P6 | 35 | M | Advanced HIV | No | 0 | 2021.01.09 | 2021.01.23 | 2021.04.22 | 104 | 4 | B.1.1.7 | Mild | Clear | Nil |
| P7 | 79 | F | Rheumatoid arthritis, | Yes | 0 | 2021.01.10 | 2021.01.20 | 2021.03.17 | 67 | 3 | B.1.1.7 | Critical | Died | Nil |
| P8 | 47 | F | Advanced HIV | No | 0 | 2021.12.01 | 2022.03.24 | 2022.06.17 | 199 | 11 | AY.4 | Mild | Clear | Remdesivir (Day 114), Regeneron (Date NA/TBC), Sotrovimab (Day 128) |
| P9 | 37 | M | Hodgkin's lymphoma | Yes | 3 | 2022.01.06 | 2022.01.22 | 2022.05.03 | 118 | 7 | BA.1.1 | Mild | Clear | Nil (TBC) |
| P10 | 67 | F | Diffuse large B-cell lymphoma, CAR-T | Yes | 3 | 2022.01.26 | 2022.01.27 | 2022.05.03 | 98 | 7 | BA.2 | Severe | Clear | Remdesivir (Day 37), Sotrovimab (Day 37), Paxlovid (Day 58) |
| P11 | 52 | F | Thymoma | No | 2 | 2022.02.09 | 2022.06.10 | 2022.06.17 | 129 | 1 | BA.1.1 | Moderate | Clear | Paxlovid (Day 128), Sotrovimab (Day 128) |
| P12 | 48 | M | Non-Hodgkin's lymphoma, CAR-T | Yes | 2 | 2022.03.20 | 2022.03.20 | 2022.10.07 | 202 | 8 | BA.2 | Mild | Clear | Sotrovimab (Day 2) |
| P13 | 52 | M | Chronic lymphoid leukaemia | Yes | 4 | 2022.03.21 | 2022.03.22 | 2022.11.14 | 239 | 1 | BA.2 | Moderate | Clear | Paxlovid (Day 5), Sotrovimab (Date NA/TBC) |
| P14 | 27 | F | Lupus | Yes | 3 | 2022.03.27 | 2022.04.30 | 2022.05.21 | 56 | 2 | BA.2 | Severe | Clear | Sotrovimab (Day 41) |
| P15 | 36 | M | Burkitt's lymphoma | Yes | 3 | 2022.04.08 | 2022.05.18 | 2022.06.02 | 56 | 2 | BA.2 | Mild | Clear | Nil (TBC) |
| P16 | 64 | M | Diffuse large B-cell lymphoma | Yes | 3 | unknown | 2022.04.19 | 2022.08.04 | 111 | 3 | BA.2 | Mild | Died | Remdesivir (Date TBC), Remdesivir (Day 17) |
| P17 | 62 | M | Diffuse large B-cell lymphoma | Yes | 4 | 2022.08.04 | 2022.07.10 | 2022.09.09 | 142 | 7 | BA.2 | Critical | Clear | Remdesivir (Days 39; 63), Sotrovimab (Days 54; 84), Paxlovid (Date TBC), Paxlovid + Remdesivir (Day 113) |
| P18 | 62 | F | Cutaneous T-cell lymphoma | No | 2 | 2022.07.12 | 2022.07.13 | 2022.09.08 | 59 | 7 | BA.5.2 | Mild | Died | Paxlovid pre-emptive (Date NA/TBC) |
| P19 | 57 | F | Simultaneous pancreas & kidney transplant | Yes | 3 | 2022.07.13 | 2022.08.02 | 2022.09.07 | 57 | 2 | BA.5.2 | Critical | Died | Sotrovimab (Day 20), Remdesivir (Date TBC) |
| P20 | 51 | M | Kidney transplant | No | 0 | 2022.09.27 | 2022.09.27 | 2022.12.01 | 66 | 5 | BA.5.2 | Mild | Unknown | Sotrovimab (Day 3), PO BD (???) (Day 3), Molnupiravir (Date NA/TBC) |
| P21 | 77 | F | MALT lymphoma | Yes | 4 | 2022.10.29 | 2022.11.15 | 2023.02.07 | 102 | 6 | BQ.1.1 | Severe | Clear | Paxlovid (Day 39), Sotrovimab (Day 40), Remdesivir (Day 40) |
| P22 | 64 | M | Diffuse large B-cell lymphoma | Yes | 3 | 2023.10.09 | 2023.10.09 | 2023.11.18 | 41 | 4 | GE.1 | Moderate | Clear | Remdesivir (Day 12), Paxlovid (Day 28) |
| P23 | 70 | M | Myeloma | No | 8 | 2023.11.23 | 2023.11.23 | 2024.01.12 | 51 | 7 | JN.1 | Critical | Died | Paxlovid (Day 27) |

### Supplemental Figure 2

#### PATIENTS 1-23

Amino-Acid Matrices representing non-synonymous mutations

Divergence Phylogeny Trees
Molecular Clock Trees

Supplemental Figure 2A

#### Patient 1

Amino-acid Matrix with NS mutations

| | 3 | 307 | 614 | 717 | 971 | 1180 |
|---|---|---|---|---|---|---|
| | V | T | D | N | G | Q |
| B.1.2 | | | G | | | |
| P1 D11 H1 | | S | G | | | |
| P1 D17 H1 | | S | G | | | |
| P1 D21 H1 | | S | G | | | |
| P1 D33 H1 | | S | G | | | |
| P1 D33 H2 | | S | G | K | | |
| P1 D33 H3 | | S | G | | | X |
| P1 D33 H4 | X | S | G | | S | |
| | 3 | 307 | 614 | 717 | 971 | 1180 |

Divergence Phylogeny Tree

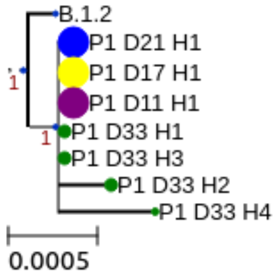


Molecular Clock Tree

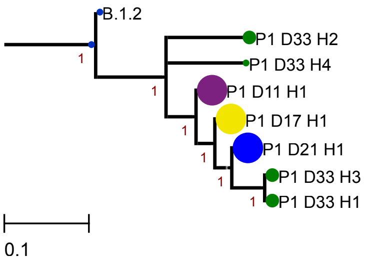


**Haplotype proportion**

- 5-20%
- 20-40%
- 40-80%
- 80-100%

**Sample (Days POS**

11, 21, 17, 33

### Supplemental Figure 2B

#### Patient 2

Amino-acid Matrix with NS mutations

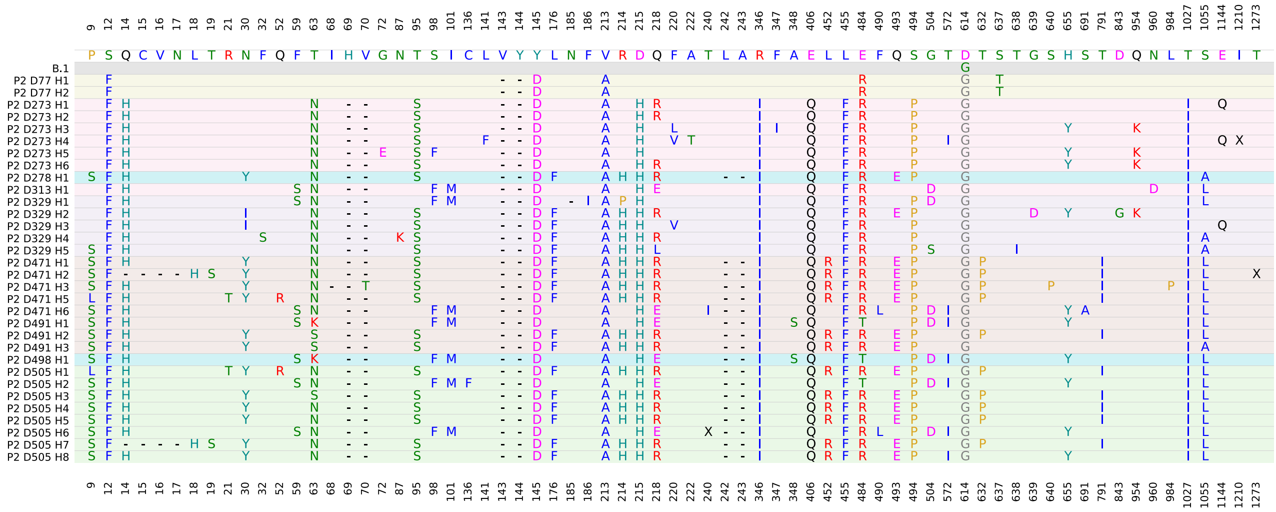

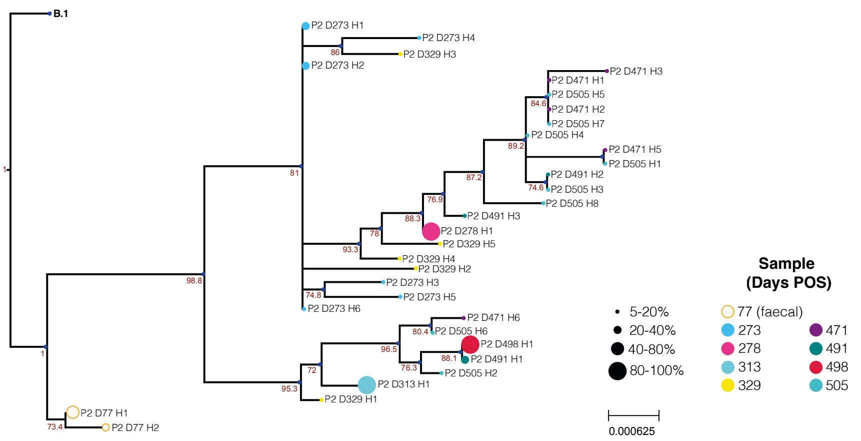


Divergence Phylogeny Tree

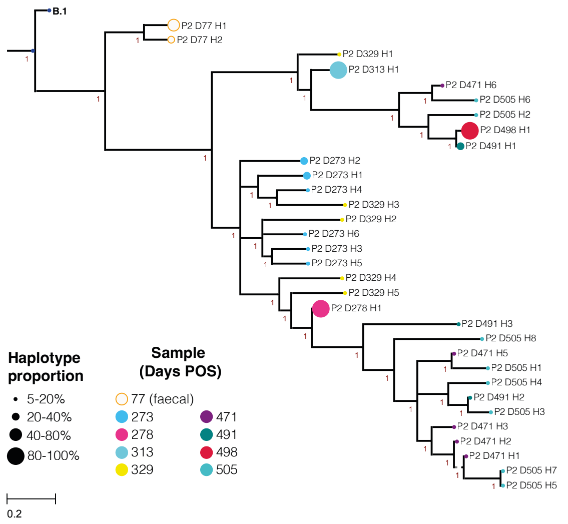


Molecular Clock Tree

### Supplemental Figure 2C

#### Patient 3

Amino-acid Matrix with NS mutations

| | 69 | 70 | 243 | 439 | 614 | 932 | 1027 | 1253 |
|---|---|---|---|---|---|---|---|---|
| | H | V | A | N | D | G | T | C |
| B.1.258 | | | | | G | | | |
| P3 D35 H1 | - | - | | K | G | D | | G |
| P3 D35 H2 | - | - | | K | G | | | G |
| P3 D35 H3 | - | - | V | K | G | | I | |
| P3 D39 H1 | - | - | | K | G | | | G |
| P3 D39 H2 | - | - | V | K | G | | I | |
| P3 D39 H3 | - | - | | K | G | | | |
| P3 D39 H4 | - | - | V | K | G | | | G |
| P3 D39 H5 | - | - | V | K | G | | I | G |
| | 69 | 70 | 243 | 439 | 614 | 932 | 1027 | 1253 |

Divergence Phylogeny Tree

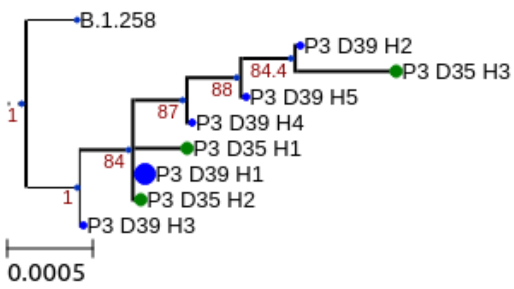


Molecular Clock Tree

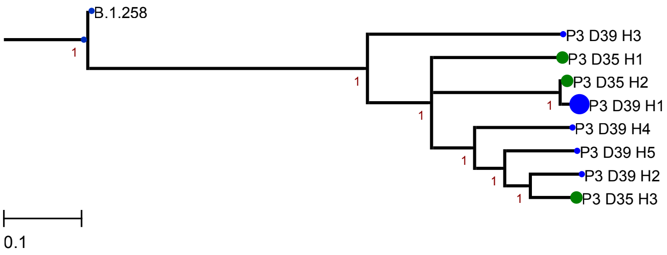


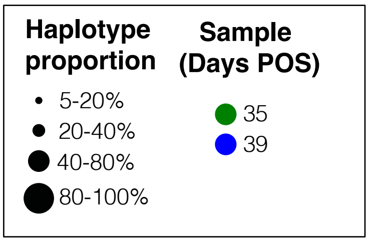

Supplemental Figure 2D

#### Patient 4

| | 42 | 307 | 355 | 509 | 614 | 1095 |
|---|---|---|---|---|---|---|
| | V | T | R | R | D | F |
| B.1.2 | | | | | G | |
| P4 D7 H1 | | S | | | G | |
| P4 D7 H2 | | S | | | G | |
| P4 D7 H3 | | S | | G | G | |
| P4 D7 H4 | A | S | K | | G | L |
| P4 D11 H1 | | S | | | G | |
| | 42 | 307 | 355 | 509 | 614 | 1095 |

Amino-acid Matrix
with NS mutations
(1st infection)

| | 90 | 222 | 583 | 614 | 1087 |
|---|---|---|---|---|---|
| | V | A | E | D | A |
| B.1.177.18 | | V | D | G | |
| P4 D0 H1 | | V | D | G | S |
| P4 D21 H1 | | V | D | G | S |
| P4 D21 H2 | A | V | D | G | S |
| P4 D31 H1 | | V | D | G | S |
| | 90 | 222 | 583 | 614 | 1087 |

Amino-acid Matrix
with NS mutations
(2nd infection)

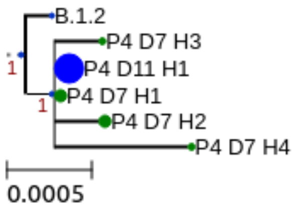


Divergence
Phylogeny Trees

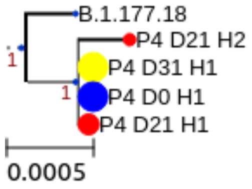


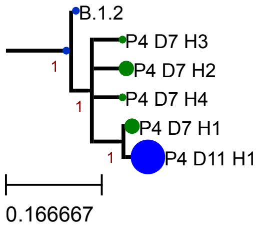


Molecular
Clock Trees

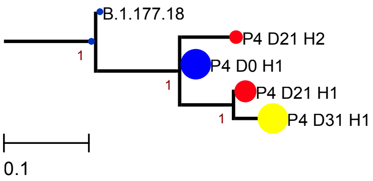


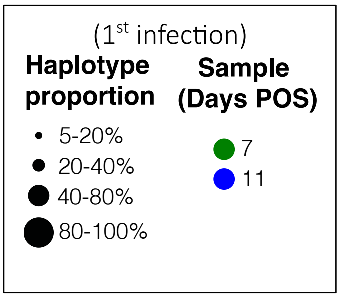


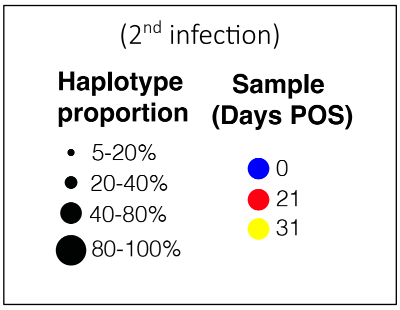

Supplemental Figure 2E

#### Patient 5

Amino-acid Matrix with NS mutations

| | 9 | 69 | 70 | 142 | 144 | 152 | 257 | 484 | 501 | 570 | 614 | 681 | 716 | 982 | 1118 |
|---|---|---|---|---|---|---|---|---|---|---|---|---|---|---|---|
| | P | H | V | G | Y | W | G | E | N | A | D | P | T | S | D |
| B.1.1.7 | | - | - | | - | | | | Y | D | G | H | I | A | H |
| P5 D12 H1 | | - | - | | - | | | | Y | D | G | H | I | A | H |
| P5 D23 H1 | | - | - | | - | | | | Y | D | G | H | I | A | H |
| P5 D39 H1 | | - | - | | - | | | | Y | D | G | H | I | A | H |
| P5 D39 H2 | L | - | - | | - | | | | Y | D | G | H | I | A | H |
| P5 D39 H3 | | - | - | | - | | S | | Y | D | G | H | I | A | H |
| P5 D48 H1 | | - | - | | - | | | | Y | D | G | H | I | A | H |
| P5 D58 H1 | | - | - | | - | L | | K | Y | D | G | H | I | A | H |
| P5 D58 H2 | | - | - | | - | | | K | Y | D | G | H | I | A | H |
| P5 D64 H1 | | - | - | - | V | | | K | Y | D | G | H | I | A | H |
| P5 D64 H2 | | - | - | - | V | | | K | Y | D | G | H | I | A | H |
| P5 D64 H3 | | - | - | | - | | | | Y | D | G | H | I | A | H |
| P5 D64 H4 | | - | - | | - | | | K | Y | D | G | H | I | A | H |
| P5 D64 H5 | | - | - | - | V | | | | Y | D | G | H | I | A | H |
| P5 D69 H1 | | - | - | - | V | | | K | Y | D | G | H | I | A | H |
| P5 D69 H2 | | - | - | - | V | | | K | Y | D | G | H | I | A | H |
| P5 D69 H3 | | - | - | | - | | | | Y | D | G | H | I | A | H |
| P5 D69 H4 | | - | - | | - | | | K | Y | D | G | H | I | A | H |
| P5 D69 H5 | | - | - | - | V | | | | Y | D | G | H | I | A | H |
| P5 D71 H1 | | - | - | | - | L | | K | Y | D | G | H | I | A | H |
| | 9 | 69 | 70 | 142 | 144 | 152 | 257 | 484 | 501 | 570 | 614 | 681 | 716 | 982 | 1118 |

Divergence Phylogeny Tree

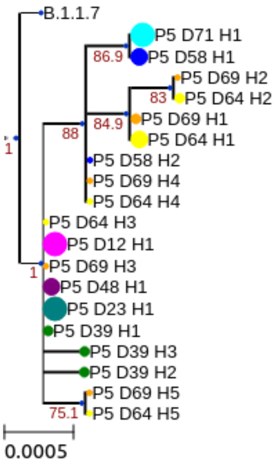


Molecular Clock Tree

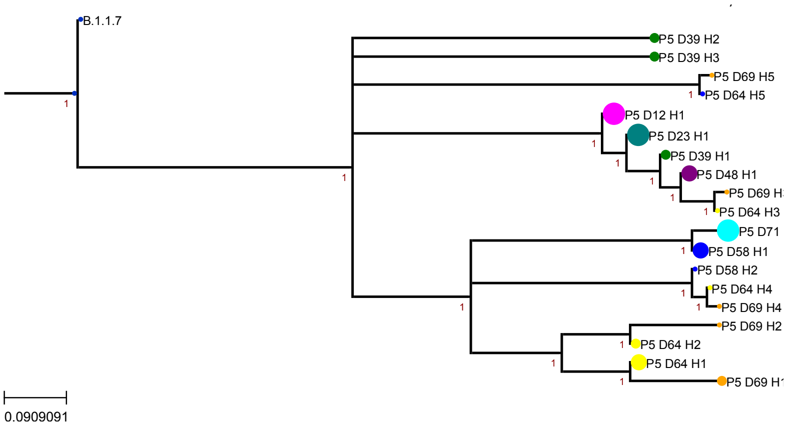


| **Haplotype proportion** | **Sample (Days POS)** | |
|---|---|---|
| • 5-20% | 12 | 58 |
| 20-40% | 23 | 64 |
| 40-80% | 39 | 69 |
| 80-100% | 48 | 71 |

Supplemental Figure 2F

Patient 6

Amino-acid Matrix with NS mutations

| | 64 | 67 | 69 | 70 | 95 | 144 | 215 | 501 | 570 | 595 | 614 | 681 | 684 | 716 | 982 | 1118 |
|---|---|---|---|---|---|---|---|---|---|---|---|---|---|---|---|---|
| | W | A | H | V | T | Y | D | N | A | V | D | P | A | T | S | D |
| B.1.1.7 | | | - | - | | - | | Y | D | | G | H | | I | A | H |
| P6 D16 H1 | | | - | - | | - | | Y | D | | G | H | | I | A | H |
| P6 D38 H1 | R | | - | - | | - | | Y | D | | G | H | V | I | A | H |
| P6 D38 H2 | R | | - | - | | - | | Y | D | | G | H | | I | A | H |
| P6 D38 H3 | | | - | - | I | - | | Y | D | | G | H | | I | A | H |
| P6 D38 H4 | | | - | - | | - | | Y | D | | G | H | | I | A | H |
| P6 D46 H1 | | V | - | - | | - | G | Y | D | A | G | H | | I | A | H |
| P6 D48 H1 | | | - | - | I | - | | Y | D | | G | H | | I | A | H |
| P6 D48 H2 | | | - | - | | - | | Y | D | | G | H | | I | A | H |
| | 64 | 67 | 69 | 70 | 95 | 144 | 215 | 501 | 570 | 595 | 614 | 681 | 684 | 716 | 982 | 1118 |

Divergence Phylogeny Tree

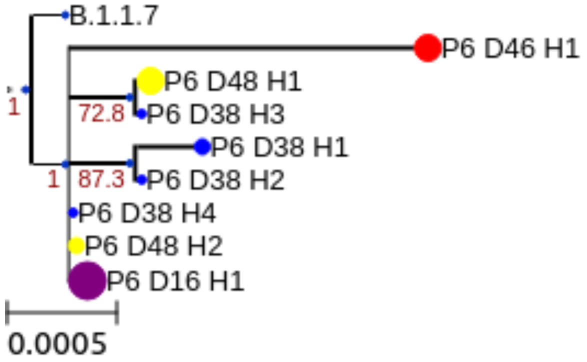


Molecular Clock Tree

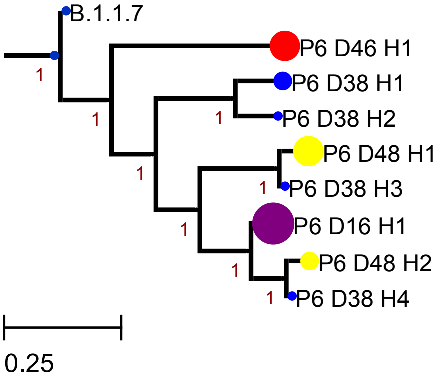


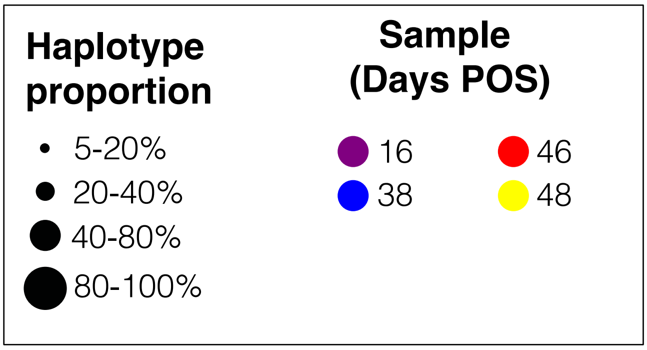

#### Supplemental Figure 2G

##### Patient 7

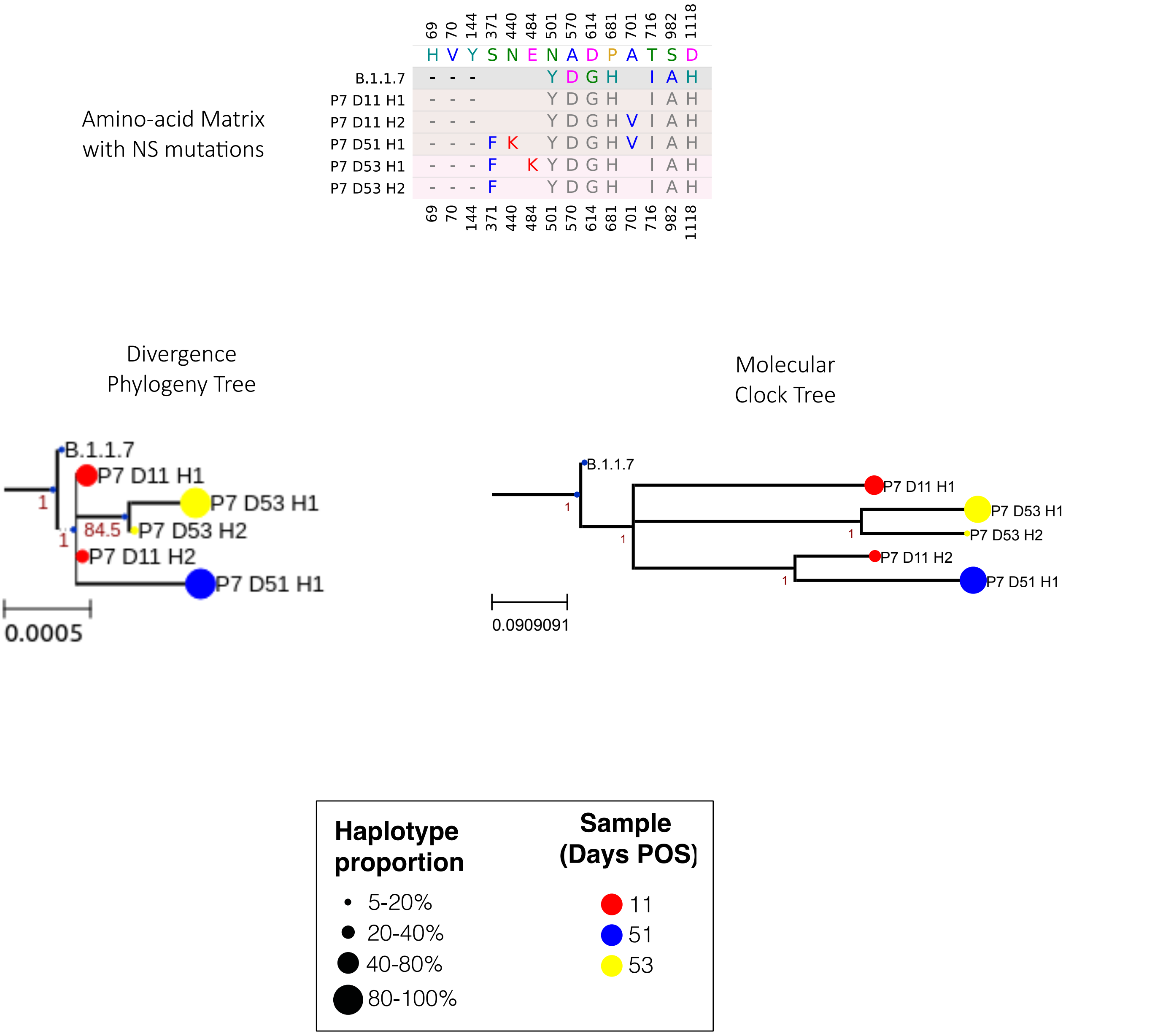

#### Supplemental Figure 2H

### Patient 8

##### Amino-acid Matrix with NS mutations

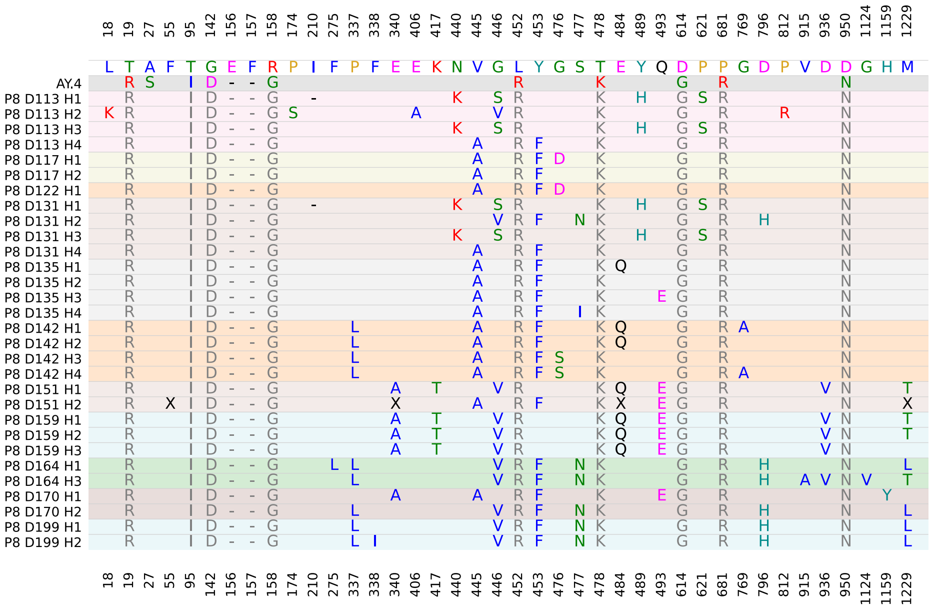


##### Divergence Phylogeny Tree

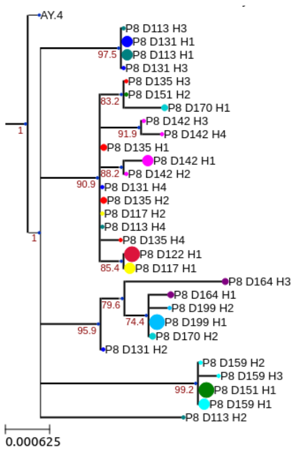


##### Molecular Clock Tree

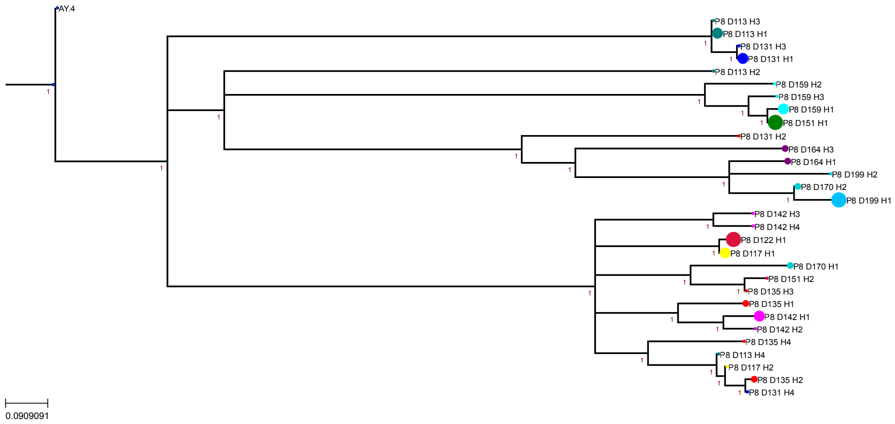


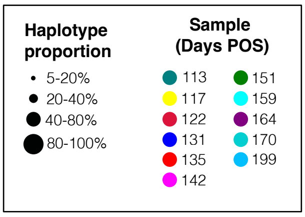

Supplemental Figure 2I

#### Patient 9

##### Amino-acid Matrix with NS mutations

| | 67 | 69 | 70 | 95 | 142 | 143 | 144 | 145 | 211 | 212 | 337 | 339 | 340 | 346 | 367 | 371 | 373 | 375 | 417 | 440 | 446 | 460 | 477 | 478 | 484 | 493 | 496 | 498 | 501 | 505 | 547 | 614 | 655 | 679 | 681 | 764 | 796 | 856 | 954 | 969 | 981 |
|---|---|---|---|---|---|---|---|---|---|---|---|---|---|---|---|---|---|---|---|---|---|---|---|---|---|---|---|---|---|---|---|---|---|---|---|---|---|---|---|---|---|
| | A | H | V | T | G | V | Y | Y | N | L | P | G | E | R | V | S | S | S | K | N | G | N | S | T | E | Q | G | Q | N | Y | T | D | H | N | P | N | D | N | Q | N | L |
| BA.1.1 | V | - | - | I | - | - | - | D | - | I | | D | | K | | | | | | K | S | | N | K | A | R | S | R | Y | H | K | G | Y | K | H | K | Y | K | H | K | F |
| P9 D16 H1 | V | - | - | I | - | - | - | D | - | I | | D | D | K | | L | P | F | N | K | S | | N | K | A | R | S | R | Y | H | K | G | Y | K | H | K | Y | K | H | K | F |
| P9 D16 H2 | V | - | - | I | - | - | - | D | - | I | S | D | | K | | L | P | F | N | K | S | | N | K | A | R | S | R | Y | H | K | G | Y | K | H | K | Y | K | H | K | F |
| P9 D16 H3 | V | - | - | I | - | - | - | D | - | I | | D | | K | | L | P | F | N | K | S | | N | K | A | R | S | R | Y | H | K | G | Y | K | H | K | Y | K | H | K | F |
| P9 D24 H1 | V | - | - | I | - | - | - | D | - | I | S | D | | K | | L | P | F | N | K | S | | N | K | A | R | S | R | Y | H | K | G | Y | K | H | K | Y | K | H | K | F |
| P9 D24 H2 | V | - | - | I | - | - | - | D | - | I | | D | | K | | L | P | F | N | K | S | | N | K | A | R | S | R | Y | H | K | G | Y | K | H | K | Y | K | H | K | F |
| P9 D24 H3 | V | - | - | I | - | - | - | D | - | I | S | D | D | K | | L | P | F | N | K | S | | N | K | A | R | S | R | Y | H | K | G | Y | K | H | K | Y | K | H | K | F |
| P9 D30 H1 | V | - | - | I | - | - | - | D | - | I | | D | D | K | | L | P | F | N | K | S | | N | K | A | R | S | R | Y | H | K | G | Y | K | H | K | Y | K | H | K | F |
| P9 D30 H2 | V | - | - | I | - | - | - | D | - | I | S | D | | K | | L | P | F | N | K | S | | N | K | A | R | S | R | Y | H | K | G | Y | K | H | K | Y | K | H | K | F |
| P9 D30 H3 | V | - | - | I | - | - | - | D | - | I | S | D | D | K | | L | P | F | N | K | S | | N | K | A | R | S | R | Y | H | K | G | Y | K | H | K | Y | K | H | K | F |
| P9 D40 H1 | V | - | - | I | - | - | - | D | - | I | S | D | | K | | L | P | F | N | K | S | K | N | K | A | R | S | R | Y | H | K | G | Y | K | H | K | Y | K | H | K | F |
| P9 D40 H2 | V | - | - | I | - | - | - | D | - | I | S | D | | K | | L | P | F | N | K | S | | N | K | A | R | S | R | Y | H | K | G | Y | K | H | K | Y | K | H | K | F |
| P9 D40 H3 | V | - | - | I | - | - | - | D | - | I | S | D | | K | F | L | P | F | N | K | S | | N | K | A | R | S | R | Y | H | K | G | Y | K | H | K | Y | K | H | K | F |
| P9 D40 H4 | V | - | - | I | - | - | - | D | - | I | | D | D | K | | L | P | F | N | K | S | | N | K | A | R | S | R | Y | H | K | G | Y | K | H | K | Y | K | H | K | F |
| P9 D43 H1 | V | - | - | I | - | - | - | D | - | I | | D | D | K | | L | P | F | N | K | S | | N | K | A | R | S | R | Y | H | K | G | Y | K | H | K | Y | K | H | K | F |
| P9 D43 H2 | V | - | - | I | - | - | - | D | - | I | S | D | | K | | L | P | F | N | K | S | K | N | K | A | R | S | R | Y | H | K | G | Y | K | H | K | Y | K | H | K | F |
| P9 D45 H1 | V | - | - | I | - | - | - | D | - | I | | D | D | K | | L | P | F | N | K | S | | N | K | A | R | S | R | Y | H | K | G | Y | K | H | K | Y | K | H | K | F |
| P9 D45 H2 | V | - | - | I | - | - | - | D | - | I | | D | D | K | | L | P | F | N | K | S | | N | K | A | R | S | R | Y | H | K | G | Y | K | H | E | Y | K | H | K | F |
| P9 D74 H1 | V | - | - | I | - | - | - | D | - | I | | D | D | K | | L | P | F | T | K | S | | N | K | A | R | S | R | Y | H | K | G | Y | K | H | K | Y | K | H | K | F |
| | 67 | 69 | 70 | 95 | 142 | 143 | 144 | 145 | 211 | 212 | 337 | 339 | 340 | 346 | 367 | 371 | 373 | 375 | 417 | 440 | 446 | 460 | 477 | 478 | 484 | 493 | 496 | 498 | 501 | 505 | 547 | 614 | 655 | 679 | 681 | 764 | 796 | 856 | 954 | 969 | 981 |

##### Divergence Phylogeny Tree

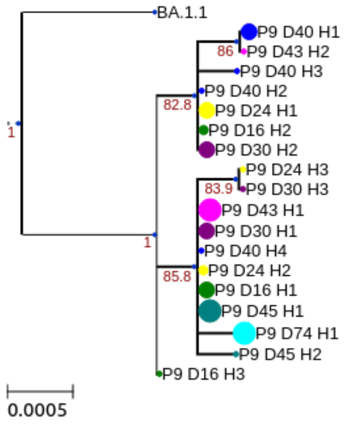


| Haplotype proportion | Sample (Days POS) | |
|---|---|---|
| • 5-20% | 16 | 43 |
| 20-40% | 24 | 45 |
| 40-80% | 30 | 74 |
| 80-100% | 40 | |

##### Molecular Clock Tree

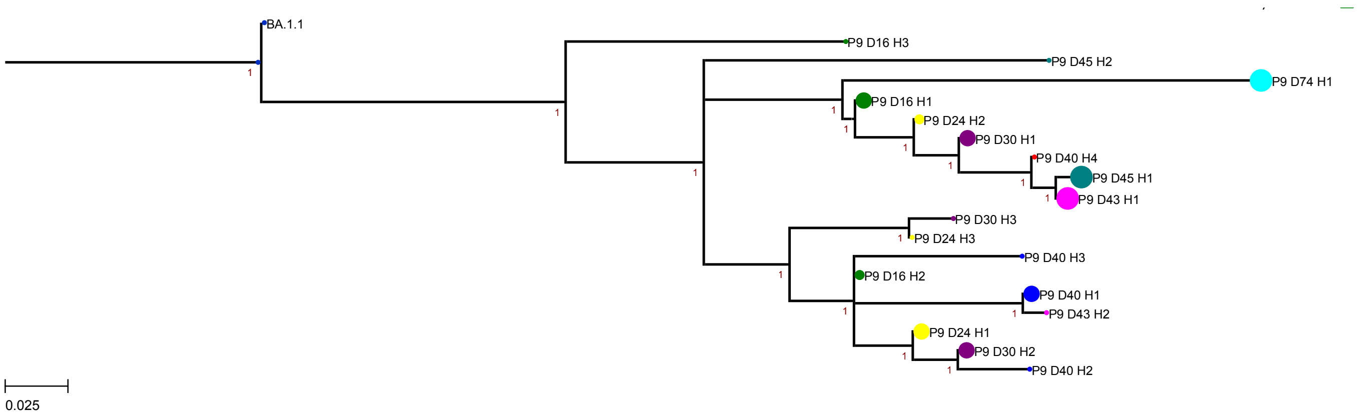

Supplemental Figure 2J

#### Patient 10

##### Amino-acid Matrix with NS mutations

| | 19 | 24 | 25 | 26 | 27 | 142 | 213 | 339 | 340 | 356 | 371 | 373 | 375 | 376 | 385 | 405 | 408 | 417 | 440 | 477 | 478 | 484 | 493 | 498 | 501 | 505 | 614 | 655 | 679 | 681 | 764 | 796 | 954 | 969 |
|---|---|---|---|---|---|---|---|---|---|---|---|---|---|---|---|---|---|---|---|---|---|---|---|---|---|---|---|---|---|---|---|---|---|---|
| | T | L | P | P | A | G | V | G | E | K | S | S | S | T | T | D | R | K | N | S | T | E | Q | Q | N | Y | D | H | N | P | N | D | Q | N |
| BA.2 | I | - | - | - | S | D | G | D | | | F | P | F | A | | N | S | N | K | N | K | A | R | R | Y | H | G | Y | K | H | K | Y | H | K |
| P10 D1 H1 | I | - | - | - | S | D | G | D | | | F | P | F | A | | N | S | N | K | N | K | A | R | R | Y | H | G | Y | K | H | K | Y | H | K |
| P10 D10 H1 | I | - | - | - | S | D | G | D | | | F | P | F | A | | N | S | N | K | N | K | A | R | R | Y | H | G | Y | K | H | K | Y | H | K |
| P10 D15 H1 | I | - | - | - | S | D | G | D | | | F | P | F | A | | N | S | N | K | N | K | A | R | R | Y | H | G | Y | K | H | K | Y | H | K |
| P10 D37 H1 | I | - | - | - | S | D | G | D | | | F | P | F | A | | N | S | N | K | N | K | A | R | R | Y | H | G | Y | K | H | K | Y | H | K |
| P10 D43 H1 | I | - | - | - | S | D | G | D | | | F | P | F | A | | N | S | N | K | N | K | A | R | R | Y | H | G | Y | K | H | K | Y | H | K |
| P10 D43 H2 | I | - | - | - | S | D | G | D | | | F | P | F | A | I | N | S | N | K | N | K | A | R | R | Y | H | G | Y | K | H | K | Y | H | K |
| P10 D43 H3 | I | - | - | - | S | D | G | D | | | F | P | F | A | I | N | S | N | K | N | K | A | R | R | Y | H | G | Y | K | H | K | Y | H | K |
| P10 D56 H1 | I | - | - | - | S | D | G | D | | | F | P | F | A | | N | S | N | K | N | K | A | R | R | Y | H | G | Y | K | H | K | Y | H | K |
| P10 D56 H2 | I | - | - | - | S | D | G | D | | T | F | P | F | | | N | S | N | K | N | K | A | R | R | Y | H | G | Y | K | H | K | Y | H | K |
| P10 D72 H1 | I | - | - | - | S | D | G | D | A | | F | P | F | A | | N | S | I | K | N | K | A | R | R | Y | H | G | Y | K | H | K | Y | H | K |

##### Divergence Phylogeny Tree

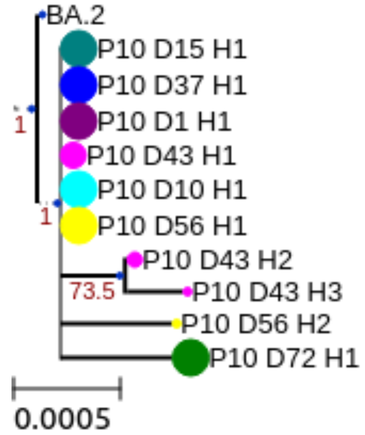


##### Molecular Clock Tree

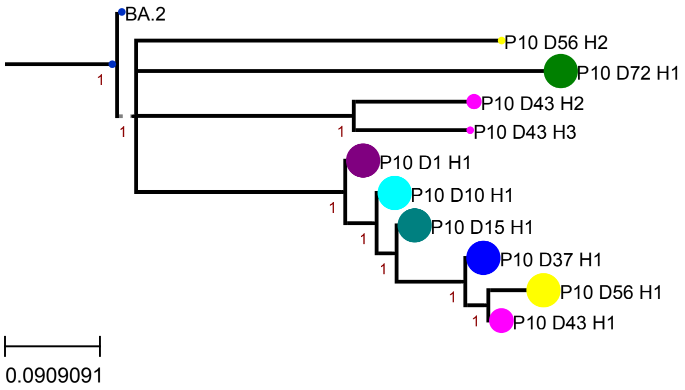


**Haplotype proportion**

- 5-20%
- 20-40%
- 40-80%
- 80-100%

**Sample (Days POS)**

1, 10, 15, 37, 43, 56, 72

Supplemental Figure 2K

#### Patient 11

##### Amino-acid Matrix with NS mutations

| | 67 | 69 | 70 | 95 | 142 | 143 | 144 | 145 | 211 | 212 | 339 | 346 | 371 | 373 | 375 | 417 | 440 | 446 | 477 | 478 | 484 | 493 | 496 | 498 | 501 | 505 | 547 | 614 | 655 | 679 | 681 | 764 | 796 | 856 | 936 | 954 | 961 | 969 | 981 | 987 |
|---|---|---|---|---|---|---|---|---|---|---|---|---|---|---|---|---|---|---|---|---|---|---|---|---|---|---|---|---|---|---|---|---|---|---|---|---|---|---|---|---|
| | A | H | V | T | G | V | Y | Y | N | L | G | R | S | S | S | K | N | G | S | T | E | Q | G | Q | N | Y | T | D | H | N | P | N | D | N | D | Q | T | N | L | V |
| BA.1.1 | V | - | - | I | - | - | - | D | - | I | D | K | | | | | K | S | N | K | A | R | S | R | Y | H | K | G | Y | K | H | K | Y | K | | H | | K | F | |
| P11 D129 H1 | V | - | - | I | - | - | - | D | - | I | D | K | L | P | F | N | K | S | N | K | A | R | S | R | Y | H | K | G | Y | K | H | K | Y | K | Y | H | | K | F | F |
| P11 D129 H2 | V | - | - | I | - | - | - | D | - | I | D | K | L | P | F | N | K | S | N | K | A | R | S | R | Y | H | K | G | Y | K | H | K | Y | K | | H | | K | F | F |
| P11 D129 H3 | V | - | - | I | - | - | - | D | - | I | D | K | L | P | F | N | K | S | N | K | A | R | S | R | Y | H | K | G | Y | K | H | K | Y | K | | H | | K | F | |
| P11 D129 H4 | V | - | - | I | - | - | - | D | - | I | D | K | I | P | F | N | K | S | N | K | A | R | S | R | Y | H | K | G | Y | K | H | K | Y | K | | H | M | K | F | F |
| P11 D129 H5 | V | - | - | I | - | - | - | D | - | I | D | K | L | P | F | N | K | S | N | K | A | R | S | R | Y | H | K | G | Y | K | H | K | Y | K | Y | H | | K | F | |
| | 67 | 69 | 70 | 95 | 142 | 143 | 144 | 145 | 211 | 212 | 339 | 346 | 371 | 373 | 375 | 417 | 440 | 446 | 477 | 478 | 484 | 493 | 496 | 498 | 501 | 505 | 547 | 614 | 655 | 679 | 681 | 764 | 796 | 856 | 936 | 954 | 961 | 969 | 981 | 987 |

##### Divergence Phylogeny Tree

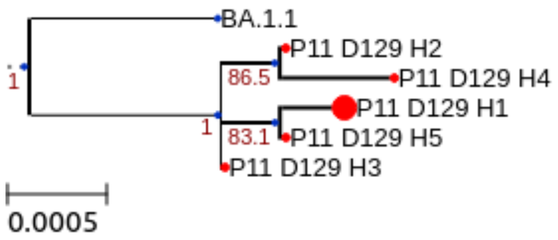


##### Molecular Clock Tree

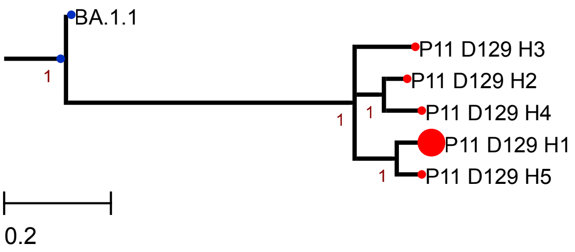


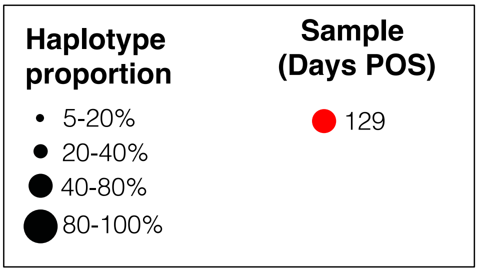

Supplemental Figure 2L

#### Patient 12

##### Amino-acid Matrix with NS mutations

| | 19 | 24 | 25 | 26 | 27 | 142 | 213 | 339 | 340 | 356 | 371 | 373 | 375 | 376 | 385 | 405 | 408 | 417 | 440 | 477 | 478 | 484 | 493 | 498 | 501 | 505 | 614 | 655 | 679 | 681 | 764 | 796 | 954 | 969 |
|---|---|---|---|---|---|---|---|---|---|---|---|---|---|---|---|---|---|---|---|---|---|---|---|---|---|---|---|---|---|---|---|---|---|---|
| | T | L | P | P | A | G | V | G | E | K | S | S | S | T | T | D | R | K | N | S | T | E | Q | Q | N | Y | D | H | N | P | N | D | Q | N |
| BA.2 | I | - | - | - | S | D | G | D | | | F | P | F | A | | N | S | N | K | N | K | A | R | R | Y | H | G | Y | K | H | K | Y | H | K |
| P12 D0 H1 | I | - | - | - | S | D | G | D | | | F | P | F | A | | N | S | N | K | N | K | A | R | R | Y | H | G | Y | K | H | K | Y | H | K |
| P12 D1 H1 | I | - | - | - | S | D | G | D | | | F | P | F | A | | N | S | N | K | N | K | A | R | R | Y | H | G | Y | K | H | K | Y | H | K |
| P12 D2 H1 | I | - | - | - | S | D | G | D | K | | F | P | F | A | | N | S | N | K | N | K | A | R | R | Y | H | G | Y | K | H | K | Y | H | K |
| P12 D24 H1 | I | - | - | - | S | D | G | D | Q | | F | P | F | A | I | N | S | N | K | N | K | A | R | R | Y | H | G | Y | K | H | K | Y | H | K |
| P12 D55 H1 | I | - | - | - | S | D | G | D | K | | F | P | F | A | | N | S | N | K | N | K | A | R | R | Y | H | G | Y | K | H | K | Y | H | K |
| P12 D55 H2 | I | - | - | - | S | D | G | D | Q | | F | P | F | A | | N | S | N | K | N | K | A | R | R | Y | H | G | Y | K | H | K | Y | H | K |
| P12 D94 H1 | I | - | - | - | S | D | G | D | Q | | F | P | F | A | I | N | S | N | K | N | K | A | R | R | Y | H | G | Y | K | H | K | Y | H | K |
| P12 D113 H1 | I | - | - | - | S | D | G | D | | R | F | P | F | A | | N | S | N | K | N | K | A | R | R | Y | H | G | Y | K | H | K | Y | H | K |
| P12 D113 H2 | I | - | - | - | S | D | G | D | | | F | P | F | A | | N | S | N | K | N | K | A | R | R | Y | H | G | Y | K | H | K | Y | H | K |
| P12 D117 H1 | I | - | - | - | S | D | G | D | | | F | P | F | A | | N | S | N | K | N | K | A | R | R | Y | H | G | Y | K | H | K | Y | H | K |
| P12 D117 H2 | I | - | - | - | S | D | G | D | | | F | P | F | A | I | N | S | N | K | N | K | A | R | R | Y | H | G | Y | K | H | K | Y | H | K |
| P12 D117 H3 | I | - | - | - | S | D | G | D | | | F | P | F | A | I | N | S | N | K | N | K | A | R | R | Y | H | G | Y | K | H | K | Y | H | K |

19 24 25 26 27 142 213 339 340 356 371 373 375 376 385 405 408 417 440 477 478 484 493 498 501 505 614 655 679 681 764 796 954 969

Divergence
Phylogeny Tree

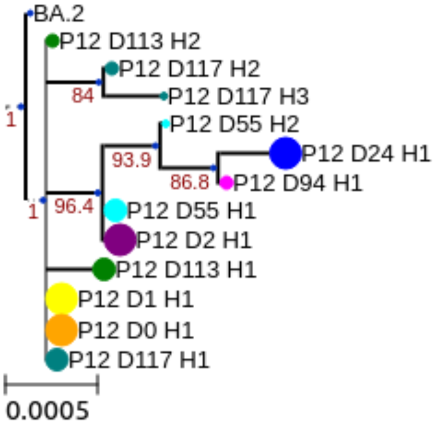


Molecular
Clock Tree

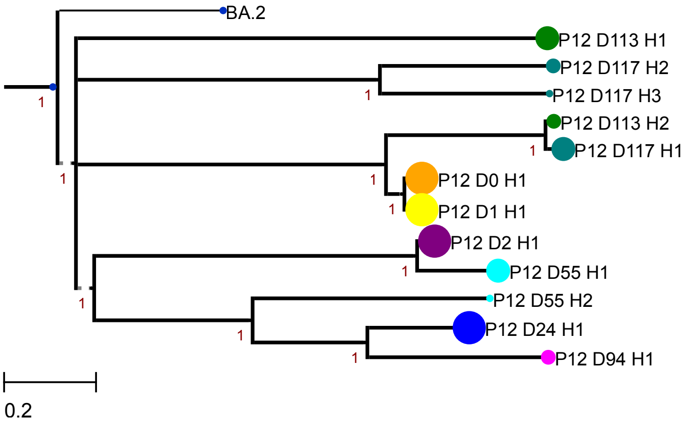


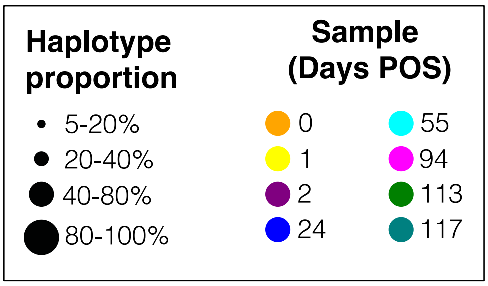

Supplemental Figure 2M

#### Patient 13

| | 19 | 24 | 25 | 26 | 27 | 142 | 213 | 299 | 308 | 339 | 371 | 373 | 375 | 376 | 385 | 405 | 408 | 417 | 440 | 477 | 478 | 484 | 493 | 498 | 501 | 505 | 614 | 655 | 679 | 681 | 764 | 796 | 944 | 954 | 969 | 1262 |
|---|---|---|---|---|---|---|---|---|---|---|---|---|---|---|---|---|---|---|---|---|---|---|---|---|---|---|---|---|---|---|---|---|---|---|---|---|
| | T | L | P | P | A | G | V | T | V | G | S | S | S | T | T | D | R | K | N | S | T | E | Q | Q | N | Y | D | H | N | P | N | D | A | Q | N | E |
| BA.2 | I | - | - | - | S | D | G | | | D | F | P | F | A | | N | S | N | K | N | K | A | R | R | Y | H | G | Y | K | H | K | Y | | H | K | |
| P13 D237 H1 | I | - | - | - | S | D | G | I | I | D | F | P | F | A | I | | S | N | K | N | K | A | R | R | Y | H | G | Y | K | H | K | Y | V | H | K | G |

TREES NOT AVAILABLE

Supplemental Figure 2N

#### Patient 14

Amino-acid Matrix with NS mutations

| | 19 | 24 | 25 | 26 | 27 | 142 | 213 | 339 | 371 | 373 | 375 | 376 | 405 | 408 | 417 | 440 | 477 | 478 | 484 | 493 | 498 | 501 | 505 | 614 | 655 | 679 | 681 | 764 | 796 | 954 | 969 |
|---|---|---|---|---|---|---|---|---|---|---|---|---|---|---|---|---|---|---|---|---|---|---|---|---|---|---|---|---|---|---|---|
| | T | L | P | P | A | G | V | G | S | S | S | T | D | R | K | N | S | T | E | Q | Q | N | Y | D | H | N | P | N | D | Q | N |
| BA.2 | I | - | - | - | S | D | G | D | F | P | F | A | N | S | N | K | N | K | A | R | R | Y | H | G | Y | K | H | K | Y | H | K |
| P14 D34 H1 | I | - | - | - | S | D | G | D | F | P | F | A | N | S | N | K | N | K | A | R | R | Y | H | G | Y | K | H | K | Y | H | K |
| P14 D34 H3 | I | - | - | - | S | D | G | D | F | P | F | A | N | S | N | K | N | K | A | R | R | Y | H | G | Y | K | H | K | H | H | K |
| P14 D39 H1 | I | - | - | - | S | D | G | D | F | P | F | A | N | S | N | K | N | K | A | R | R | Y | H | G | Y | K | H | K | Y | H | K |
| | 19 | 24 | 25 | 26 | 27 | 142 | 213 | 339 | 371 | 373 | 375 | 376 | 405 | 408 | 417 | 440 | 477 | 478 | 484 | 493 | 498 | 501 | 505 | 614 | 655 | 679 | 681 | 764 | 796 | 954 | 969 |

Divergence
Phylogeny Tree

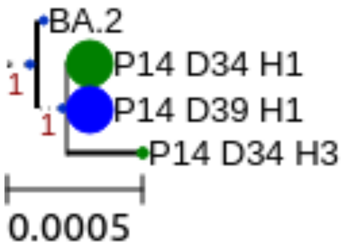


Molecular
Clock Tree

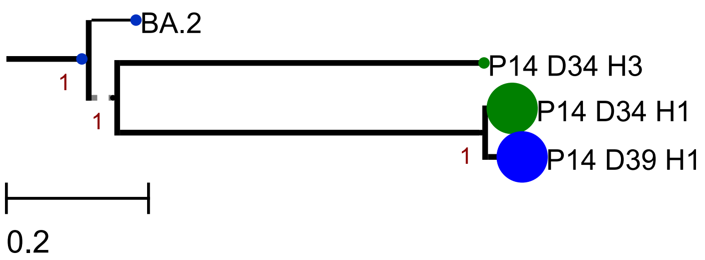


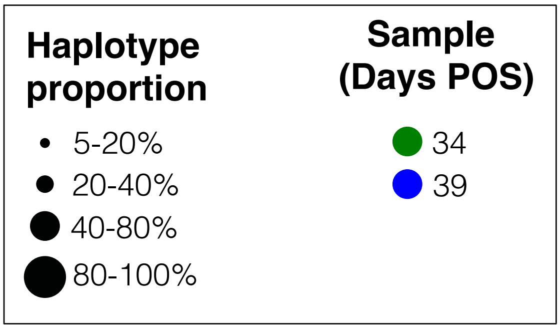

Supplemental Figure 2O

#### Patient 15

Amino-acid Matrix with NS mutations

| | 19 | 24 | 25 | 26 | 27 | 142 | 213 | 248 | 339 | 371 | 373 | 375 | 376 | 405 | 408 | 417 | 440 | 477 | 478 | 484 | 493 | 498 | 501 | 505 | 614 | 655 | 679 | 681 | 748 | 764 | 796 | 954 | 969 |
|---|---|---|---|---|---|---|---|---|---|---|---|---|---|---|---|---|---|---|---|---|---|---|---|---|---|---|---|---|---|---|---|---|---|
| | T | L | P | P | A | G | V | Y | G | S | S | S | T | D | R | K | N | S | T | E | Q | Q | N | Y | D | H | N | P | E | N | D | Q | N |
| BA.2 | I | - | - | - | S | D | G | | D | F | P | F | A | N | S | N | K | N | K | A | R | R | Y | H | G | Y | K | H | | K | Y | H | K |
| P15 D40 H1 | I | - | - | - | S | D | G | H | D | F | P | F | A | N | S | N | K | N | K | A | R | R | Y | H | G | Y | K | H | Q | K | Y | H | K |
| P15 D44 H1 | I | - | - | - | S | D | G | H | D | F | P | F | A | N | S | N | K | N | K | A | R | R | Y | H | G | Y | K | H | Q | K | Y | H | K |

Divergence Phylogeny Tree

Molecular Clock Tree

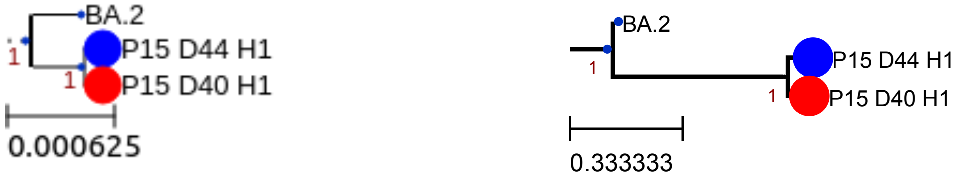


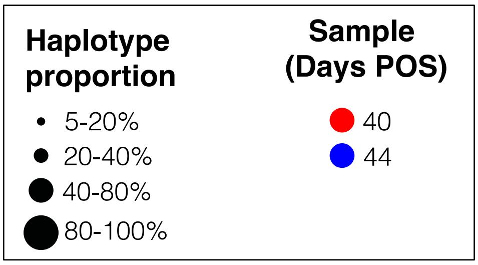

Supplemental Figure 2P

#### Patient 16

Amino-acid Matrix with NS mutations

| | 19 | 24 | 25 | 26 | 27 | 109 | 142 | 213 | 339 | 371 | 373 | 375 | 376 | 405 | 408 | 417 | 440 | 477 | 478 | 484 | 493 | 498 | 501 | 505 | 515 | 614 | 655 | 679 | 681 | 764 | 796 | 954 | 969 | 1264 |
|---|---|---|---|---|---|---|---|---|---|---|---|---|---|---|---|---|---|---|---|---|---|---|---|---|---|---|---|---|---|---|---|---|---|---|
| | T | L | P | P | A | T | G | V | G | S | S | S | T | D | R | K | N | S | T | E | Q | Q | N | Y | F | D | H | N | P | N | D | Q | N | V |
| BA.2 | I | - | - | - | S | | D | G | D | F | P | F | A | N | S | N | K | N | K | A | R | R | Y | H | | G | Y | K | H | K | Y | H | K | |
| P16 D0 H1 | I | - | - | - | S | | D | G | D | F | P | F | A | N | S | N | K | N | K | A | R | R | Y | H | | G | Y | K | H | K | Y | H | K | |
| P16 D0 H2 | I | - | - | - | S | | D | G | D | F | P | F | A | N | S | N | K | N | K | A | R | R | Y | H | S | G | Y | K | H | K | Y | H | K | |
| P16 D0 H3 | I | - | - | - | S | | D | G | D | F | P | F | A | N | S | N | K | N | K | A | R | R | Y | H | | G | Y | K | H | K | Y | H | K | A |
| P16 D0 H4 | I | - | - | - | S | | D | G | D | F | P | F | A | N | S | N | K | N | K | A | R | R | Y | H | S | G | Y | K | H | K | Y | H | K | |
| P16 D5 H1 | I | - | - | - | S | | D | G | D | F | P | F | A | N | S | N | K | N | K | A | R | R | Y | H | | G | Y | K | H | K | Y | H | K | |
| P16 D6 H1 | I | - | - | - | S | A | D | G | D | F | P | F | A | N | S | N | K | N | K | A | R | R | Y | H | | G | Y | K | H | K | Y | H | K | |
| | 19 | 24 | 25 | 26 | 27 | 109 | 142 | 213 | 339 | 371 | 373 | 375 | 376 | 405 | 408 | 417 | 440 | 477 | 478 | 484 | 493 | 498 | 501 | 505 | 515 | 614 | 655 | 679 | 681 | 764 | 796 | 954 | 969 | 1264 |

Divergence Phylogeny Tree

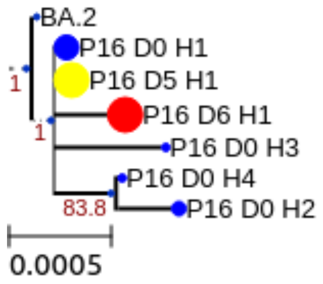


Molecular Clock Tree

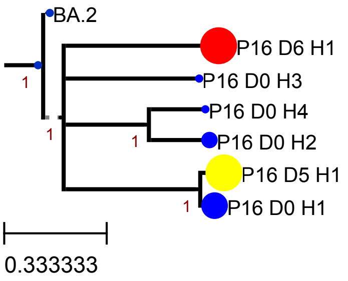


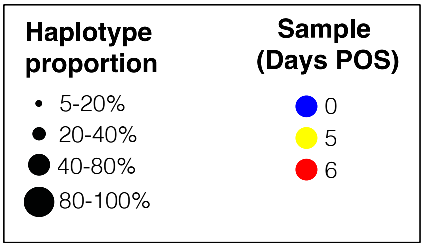

#### Supplemental Figure 2Q

##### Patient 17

Amino-acid Matrix with NS mutations

| | 19 | 24 | 25 | 26 | 27 | 142 | 213 | 334 | 337 | 339 | 340 | 344 | 346 | 356 | 357 | 370 | 371 | 373 | 375 | 376 | 405 | 408 | 417 | 440 | 477 | 478 | 484 | 493 | 498 | 501 | 504 | 505 | 614 | 655 | 679 | 681 | 764 | 785 | 796 | 954 | 969 | 1003 |
|---|---|---|---|---|---|---|---|---|---|---|---|---|---|---|---|---|---|---|---|---|---|---|---|---|---|---|---|---|---|---|---|---|---|---|---|---|---|---|---|---|---|---|
| | T | L | P | P | A | G | V | N | P | G | E | A | R | K | R | N | S | S | S | T | D | R | K | N | S | T | E | Q | Q | N | G | Y | D | H | N | P | N | V | D | Q | N | S |
| BA.2 | I | - | - | - | S | D | G | | | D | | | | | | | F | P | F | A | N | S | N | K | N | K | A | R | R | Y | | H | G | Y | K | H | K | | Y | H | K | |
| P17 D80 H1 | I | - | - | - | S | D | G | | | D | | | | R | | | F | P | F | A | N | S | N | K | N | K | A | R | R | Y | | H | G | Y | K | H | K | | Y | H | K | |
| P17 D81 H1 | I | - | - | - | S | D | G | | | D | | | | R | | | F | P | F | A | N | S | N | K | N | K | A | R | R | Y | | H | G | Y | K | H | K | | Y | H | K | |
| P17 D95 H1 | I | - | - | - | S | D | G | | | D | | | I | R | | | F | P | F | A | N | S | N | K | N | K | A | R | R | Y | | H | G | Y | K | H | K | | Y | H | K | |
| P17 D95 H2 | I | - | - | - | S | D | G | | | D | | S | | R | | | F | P | F | A | N | S | N | K | N | K | A | R | R | Y | | H | G | Y | K | H | K | | Y | H | K | |
| P17 D95 H3 | I | - | - | - | S | D | G | | | D | | | | R | | | F | P | F | A | N | S | N | K | N | K | A | R | R | Y | | H | G | Y | K | H | K | | Y | H | K | |
| P17 D95 H4 | I | - | - | - | S | D | G | | | D | | | | R | | | F | P | F | A | N | S | N | K | N | K | A | R | R | Y | | H | G | Y | K | H | K | | Y | H | K | |
| P17 D95 H5 | I | - | - | - | S | D | G | | | D | | S | | R | | | F | P | F | A | N | S | N | K | N | K | A | R | R | Y | | H | G | Y | K | H | K | | Y | H | K | |
| P17 D95 H6 | I | - | - | - | S | D | G | | | D | | | I | R | | | F | P | F | A | N | S | N | K | N | K | A | R | R | Y | | H | G | Y | K | H | K | | Y | H | K | |
| P17 D104 H1 | I | - | - | - | S | D | G | | S | D | | | | | | K | F | P | F | A | N | S | N | K | N | K | A | R | R | Y | D | H | G | Y | K | H | K | | Y | H | K | |
| P17 D104 H2 | I | - | - | - | S | D | G | | | D | D | | | | K | | F | P | F | A | N | S | N | K | N | K | A | R | R | Y | D | H | G | Y | K | H | K | G | Y | H | K | |
| P17 D104 H3 | I | - | - | - | S | D | G | | S | D | | | | | | K | F | P | F | A | N | S | N | K | N | K | A | R | R | Y | D | H | G | Y | K | H | K | G | Y | H | K | |
| P17 D104 H4 | I | - | - | - | S | D | G | | | D | D | | | | K | | F | P | F | A | N | S | N | K | N | K | A | R | R | Y | D | H | G | Y | K | H | K | | Y | H | K | |
| P17 D105 H1 | I | - | - | - | S | D | G | | | D | | | I | R | | | F | P | F | A | N | S | N | K | N | K | A | R | R | Y | | H | G | Y | K | H | K | | Y | H | K | |
| P17 D108 H1 | I | - | - | - | S | D | G | | | D | | | I | R | | | F | P | F | A | N | S | N | K | N | K | A | R | R | Y | | H | G | Y | K | H | K | | Y | H | K | |
| P17 D108 H2 | I | - | - | - | S | D | G | K | | D | | | I | R | | | F | P | F | A | N | S | N | K | N | K | A | R | R | Y | | H | G | Y | K | H | K | | Y | H | K | |
| P17 D108 H3 | I | - | - | - | S | D | G | | | D | | | | R | | | F | P | F | A | N | S | N | K | N | K | A | R | R | Y | | H | G | Y | K | H | K | | Y | H | K | |
| P17 D114 H1 | I | - | - | - | S | D | G | K | | D | | | I | R | | | F | P | F | A | N | S | N | K | N | K | A | R | R | Y | | H | G | Y | K | H | K | | Y | H | K | |
| P17 D114 H2 | I | - | - | - | S | D | G | | | D | | | I | R | | | F | P | F | A | N | S | N | K | N | K | A | R | R | Y | | H | G | Y | K | H | K | | Y | H | K | I |
| P17 D114 H3 | I | - | - | - | S | D | G | K | | D | | | I | R | | | F | P | F | A | N | S | N | K | N | K | A | R | R | Y | | H | G | Y | K | H | K | | Y | H | K | |
| P17 D114 H4 | I | - | - | - | S | D | G | | | D | A | | | R | | | F | P | F | A | N | S | N | K | N | K | A | R | R | Y | | H | G | Y | K | H | K | | Y | H | K | |

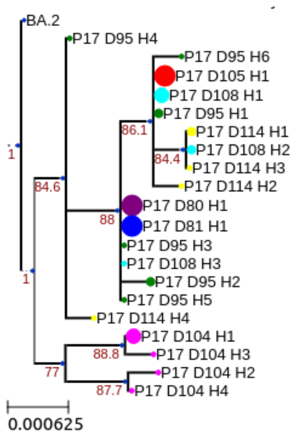


Divergence Phylogeny Tree

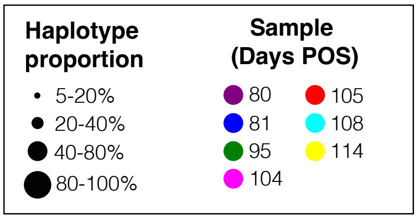


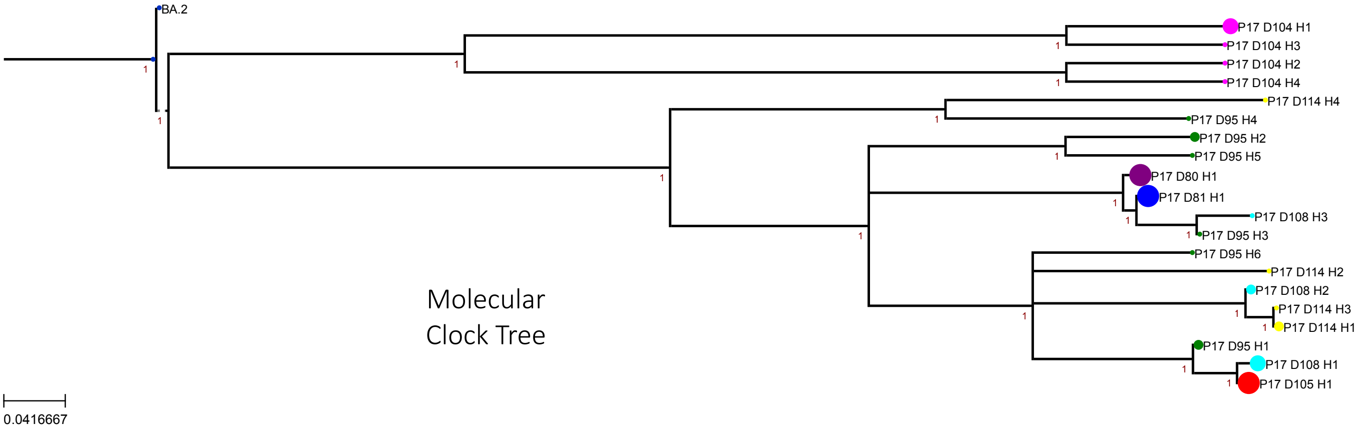


Molecular Clock Tree

Supplemental Figure 2R

#### Patient 18

Amino-acid Matrix with NS mutations

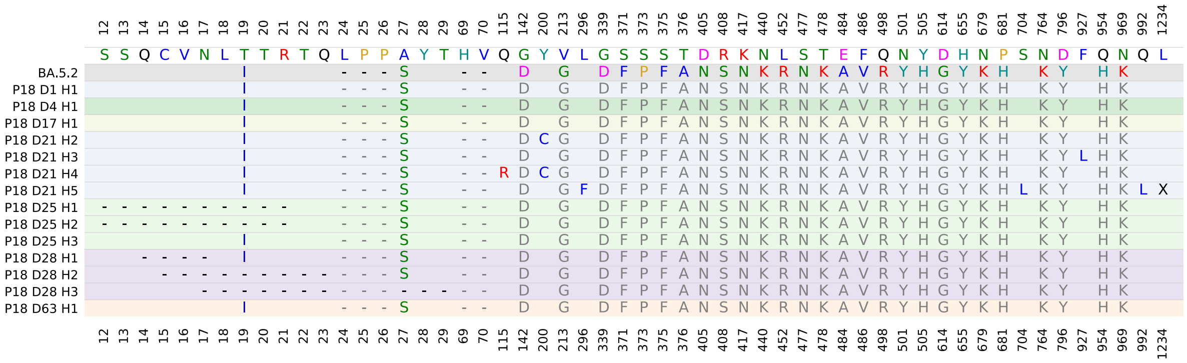


Divergence Phylogeny Tree

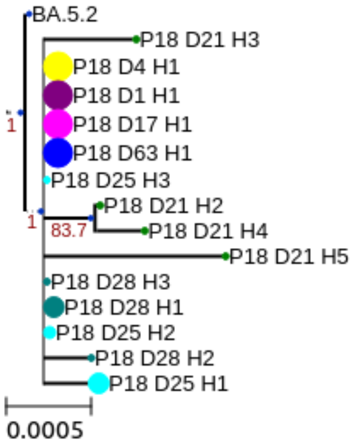


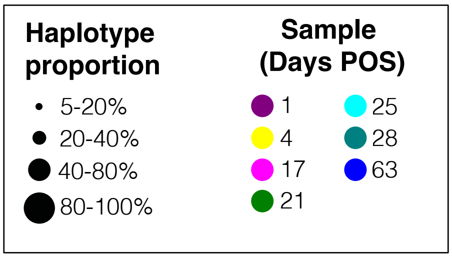


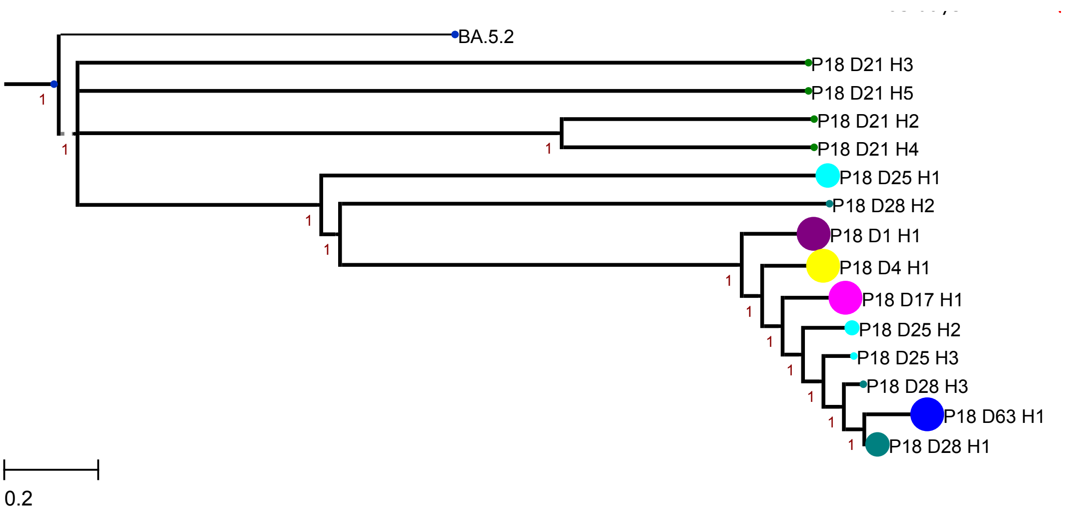


Molecular Clock Tree

Supplemental Figure 2S

#### Patient 19

Amino-acid Matrix with NS mutations

| | 19 | 24 | 25 | 26 | 27 | 64 | 69 | 70 | 139 | 140 | 141 | 142 | 143 | 144 | 145 | 173 | 184 | 210 | 213 |
|---|---|---|---|---|---|---|---|---|---|---|---|---|---|---|---|---|---|---|---|
| | T | L | P | P | A | W | H | V | P | F | L | G | V | Y | Y | Q | G | I | V |
| BA.5.2 | I | - | - | - | S | | - | - | | | | D | | | | | | | G |
| P19 D20 H1 | I | - | - | - | S | | - | - | | | | D | | | | | | | G |
| P19 D56 H1 | I | - | - | - | S | L | - | - | - | - | - | - | - | - | - | R | V | T | G |

| | 339 | 340 | 371 | 373 | 375 | 376 | 405 | 408 | 417 | 435 | 440 | 452 | 477 | 478 | 484 | 486 | 498 | 501 | 505 | 570 | 574 | 614 | 655 | 679 | 681 | 764 | 796 | 954 | 969 |
|---|---|---|---|---|---|---|---|---|---|---|---|---|---|---|---|---|---|---|---|---|---|---|---|---|---|---|---|---|---|
| | G | E | S | S | S | T | D | R | K | A | N | L | S | T | E | F | Q | N | Y | A | D | D | H | N | P | N | D | Q | N |
| BA.5.2 | D | | F | P | F | A | N | S | N | | K | R | N | K | A | V | R | Y | H | | | G | Y | K | H | K | Y | H | K |
| P19 D20 H1 | D | | F | P | F | A | N | S | N | | K | R | N | K | A | V | R | Y | H | | | G | Y | K | H | K | Y | H | K |
| P19 D56 H1 | D | K | F | P | F | A | N | S | N | T | K | R | N | K | V | V | R | Y | | T | N | G | Y | K | H | K | Y | H | K |

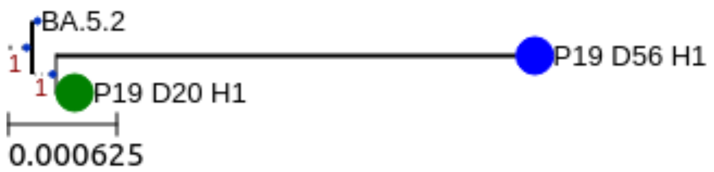


Divergence Phylogeny Tree

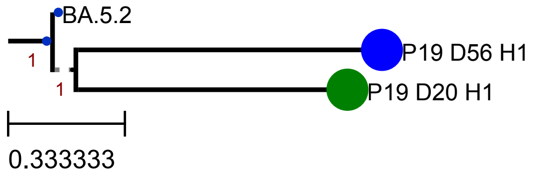


Molecular Clock Tree

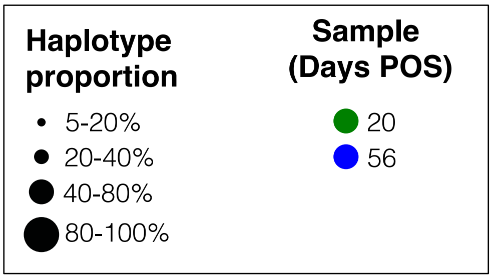

Supplemental Figure 2T

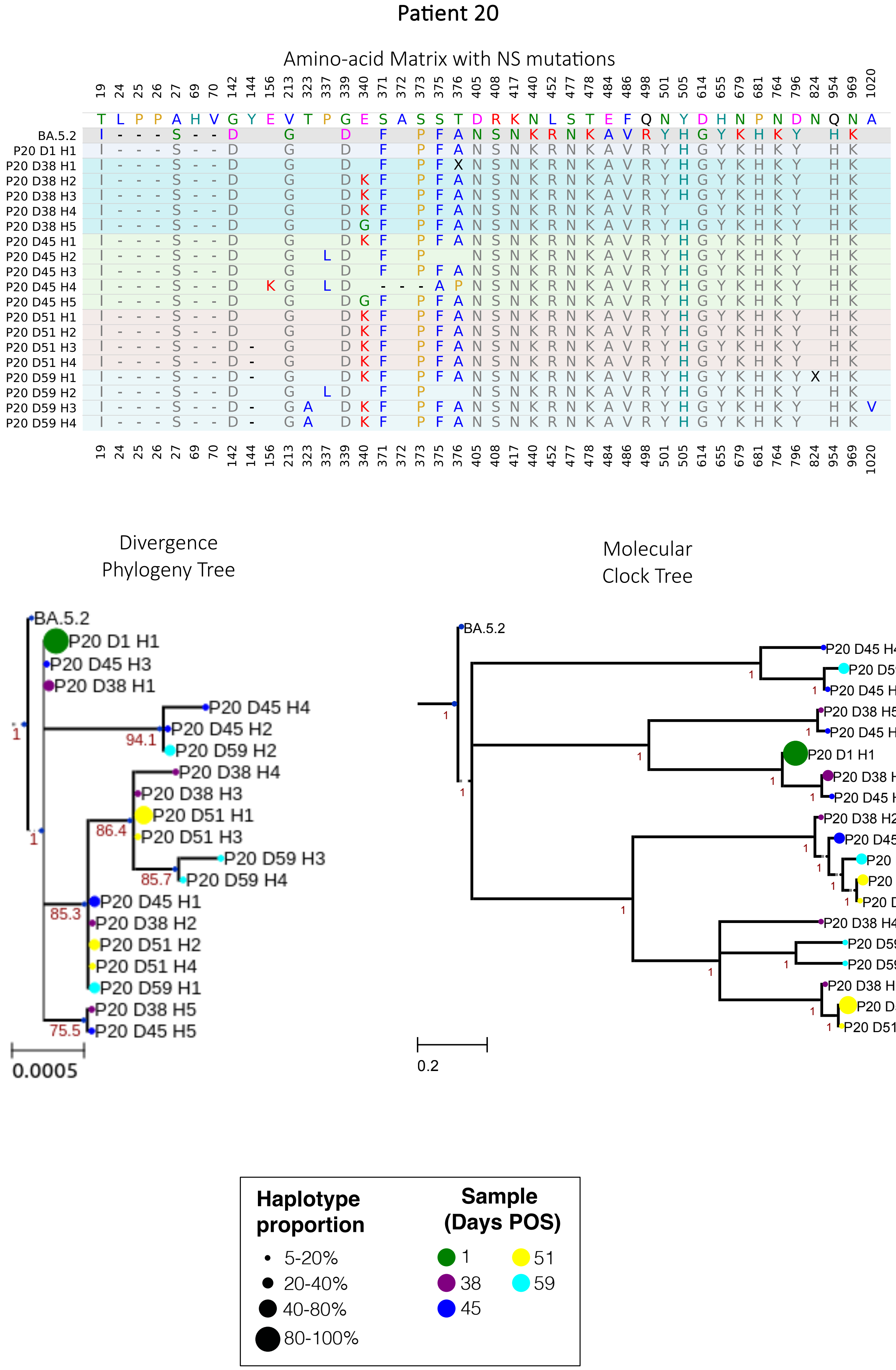

Supplemental Figure 2U

#### Patient 21

##### Amino-acid Matrix with NS mutations

| | 19 | 24 | 25 | 26 | 27 | 68 | 69 | 70 | 105 | 142 | 213 | 330 | 339 | 346 | 371 | 373 | 375 | 376 | 405 | 408 |
|---|---|---|---|---|---|---|---|---|---|---|---|---|---|---|---|---|---|---|---|---|
| | T | L | P | P | A | I | H | V | I | G | V | P | G | R | S | S | S | T | D | R |
| BQ.1.1 | I | - | - | - | S | | - | - | | D | G | | D | T | F | P | F | A | N | S |
| P21 D17 H2 | I | - | - | - | S | | - | - | | D | G | S | D | T | F | P | F | A | N | S |
| P21 D17 H3 | I | - | - | - | S | | - | - | X | D | G | | D | T | F | P | F | A | N | S |
| P21 D23 H1 | I | - | - | - | S | | - | - | | D | G | S | D | T | F | P | F | A | N | S |
| P21 D26 H1 | I | - | - | - | S | | - | - | | D | G | S | D | T | F | P | F | A | N | S |
| P21 D29 H1 | I | - | - | - | S | | - | - | | D | G | S | D | T | F | P | F | A | N | S |
| P21 D29 H2 | I | - | - | - | S | | - | - | | D | G | S | D | T | F | P | F | A | N | S |
| P21 D29 H4 | I | - | - | - | S | - | - | T | | D | G | S | D | T | F | P | F | A | N | S |
| P21 D33 H1 | I | - | - | - | S | | - | - | | D | G | S | D | T | F | P | F | A | N | S |
| P21 D33 H2 | I | - | - | - | S | | - | - | | D | G | | D | T | F | P | F | A | N | S |
| P21 D42 H1 | I | - | - | - | S | | - | - | | D | G | S | D | T | F | P | F | A | N | S |
| P21 D42 H2 | I | - | - | - | S | | - | - | | D | G | | D | T | F | P | F | A | N | S |

| | 417 | 440 | 444 | 452 | 460 | 477 | 478 | 484 | 486 | 498 | 501 | 505 | 614 | 655 | 679 | 681 | 752 | 764 | 796 | 954 | 969 | 1072 | 1083 | 1103 |
|---|---|---|---|---|---|---|---|---|---|---|---|---|---|---|---|---|---|---|---|---|---|---|---|---|
| | K | N | K | L | N | S | T | E | F | Q | N | Y | D | H | N | P | L | N | D | Q | N | E | H | F |
| BQ.1.1 | N | K | T | R | K | N | K | A | V | R | Y | H | G | Y | K | H | | K | Y | H | K | | | |
| P21 D17 H2 | N | K | T | R | K | N | K | A | V | R | Y | H | G | Y | K | H | | K | Y | H | K | | | |
| P21 D17 H3 | N | K | T | R | K | N | K | A | V | R | Y | H | G | Y | K | H | | K | Y | H | K | X | | |
| P21 D23 H1 | N | K | T | R | K | N | K | A | V | R | Y | H | G | Y | K | H | | K | Y | H | K | | | C |
| P21 D26 H1 | N | K | T | R | K | N | K | A | V | R | Y | H | G | Y | K | H | | K | Y | H | K | | | |
| P21 D29 H1 | N | K | T | R | K | N | K | A | V | R | Y | H | G | Y | K | H | P | K | Y | H | K | | | |
| P21 D29 H2 | N | K | T | R | K | N | K | A | V | R | Y | H | G | Y | K | H | | K | Y | H | K | | | |
| P21 D29 H4 | N | K | T | R | K | N | K | A | V | R | Y | H | G | Y | K | H | | K | Y | H | K | | L | |
| P21 D33 H1 | N | K | T | R | K | N | K | A | V | R | Y | H | G | Y | K | H | | K | Y | H | K | | | |
| P21 D33 H2 | N | K | T | R | K | N | K | A | V | R | Y | H | G | Y | K | H | | K | Y | H | K | | | |
| P21 D42 H1 | N | K | T | R | K | N | K | A | V | R | Y | H | G | Y | K | H | | K | Y | H | K | | | |
| P21 D42 H2 | N | K | T | R | K | N | K | A | V | R | Y | H | G | Y | K | H | | K | Y | H | K | | | |

Divergence Phylogeny Tree

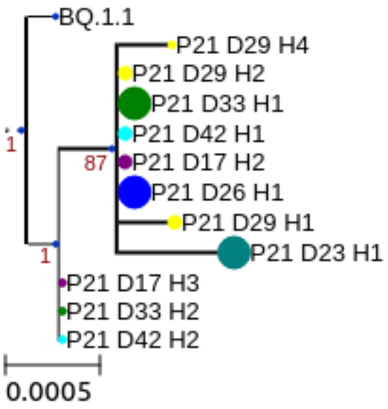


Molecular Clock Tree

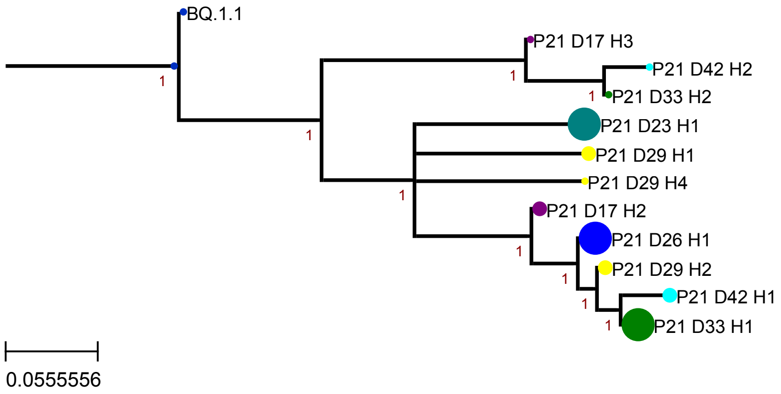


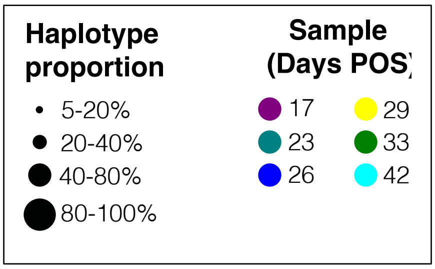

Supplemental Figure 2V

#### Patient 22

##### Amino-acid Matrix with NS mutations

| | 19 | 24 | 25 | 26 | 27 | 83 | 142 | 144 | 146 | 183 | 185 | 186 | 213 | 253 | 339 | 346 | 367 | 368 | 371 | 373 | 375 | 376 | 405 | 408 | 417 | 440 | 445 | 446 | 460 | 477 | 478 | 484 | 486 | 490 | 498 | 501 | 505 | 513 | 521 | 614 | 655 | 679 | 681 | 764 | 796 | 954 | 969 |
|---|---|---|---|---|---|---|---|---|---|---|---|---|---|---|---|---|---|---|---|---|---|---|---|---|---|---|---|---|---|---|---|---|---|---|---|---|---|---|---|---|---|---|---|---|---|---|---|
| | T | L | P | P | A | V | G | Y | H | Q | N | F | V | D | G | R | V | L | S | S | S | T | D | R | K | N | V | G | N | S | T | E | F | F | Q | N | Y | L | P | D | H | N | P | N | D | Q | N |
| GE.1 | I | - | - | - | S | A | D | - | Q | E | - | I | E | G | H | T | | I | F | P | F | A | N | S | N | K | P | S | K | N | R | A | P | S | R | Y | H | | S | G | Y | K | H | K | Y | H | K |
| P22 D0 H1 | I | - | - | - | S | A | D | - | Q | E | - | I | E | G | H | T | F | I | F | P | F | A | N | S | N | K | P | S | K | N | R | A | P | S | R | Y | H | | S | G | Y | K | H | K | Y | H | K |
| P22 D3 H1 | I | - | - | - | S | A | D | - | Q | E | - | I | E | G | H | T | F | I | F | P | F | A | N | S | N | K | P | S | K | N | R | A | P | S | R | Y | H | F | S | G | Y | K | H | K | Y | H | K |
| P22 D3 H2 | I | - | - | - | S | A | D | - | Q | E | - | I | E | G | H | T | F | I | F | P | F | A | N | S | N | K | P | S | K | N | R | A | P | S | R | Y | H | | S | G | Y | K | H | K | Y | H | K |
| P22 D10 H1 | I | - | - | - | S | A | D | - | Q | E | - | I | E | G | H | T | F | I | F | P | F | A | N | S | N | K | P | S | K | N | R | A | P | S | R | Y | H | F | S | G | Y | K | H | K | Y | H | K |
| P22 D10 H2 | I | - | - | - | S | A | D | - | Q | E | - | I | E | G | H | T | | I | F | P | F | A | N | S | N | K | P | S | K | N | R | A | P | S | R | Y | H | | S | G | Y | K | H | K | Y | H | K |
| P22 D10 H3 | I | - | - | - | S | A | D | - | Q | E | - | I | E | G | H | T | | I | F | P | F | A | N | S | N | K | P | S | K | N | R | A | P | S | R | Y | H | F | S | G | Y | K | H | K | Y | H | K |
| P22 D10 H4 | I | - | - | - | S | A | D | - | Q | E | - | I | E | G | H | T | F | I | F | P | F | A | N | S | N | K | P | S | K | N | R | A | P | S | R | Y | H | | S | G | Y | K | H | K | Y | H | K |
| P22 D21 H1 | I | - | - | - | S | A | D | - | Q | E | - | I | E | G | H | T | | I | F | P | F | A | N | S | N | K | P | S | K | N | R | A | P | S | R | Y | H | | S | G | Y | K | H | K | Y | H | K |
| P22 D21 H2 | I | - | - | - | S | A | D | - | Q | E | - | I | E | G | H | T | F | I | F | P | F | A | N | S | N | K | P | S | K | N | R | A | P | S | R | Y | H | F | S | G | Y | K | H | K | Y | H | K |
| | 19 | 24 | 25 | 26 | 27 | 83 | 142 | 144 | 146 | 183 | 185 | 186 | 213 | 253 | 339 | 346 | 367 | 368 | 371 | 373 | 375 | 376 | 405 | 408 | 417 | 440 | 445 | 446 | 460 | 477 | 478 | 484 | 486 | 490 | 498 | 501 | 505 | 513 | 521 | 614 | 655 | 679 | 681 | 764 | 796 | 954 | 969 |

##### Divergence Phylogeny Tree

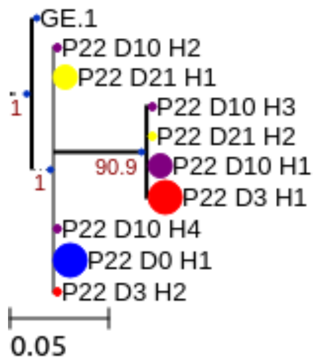


##### Molecular Clock Tree

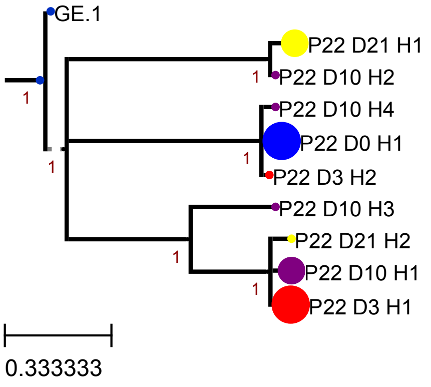


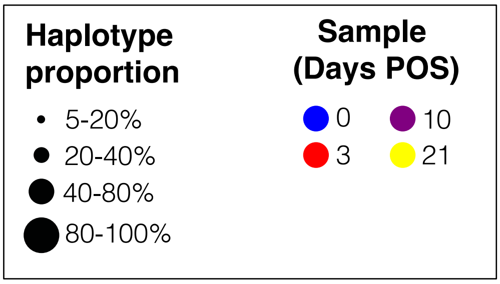

#### Supplemental Figure 2W

### Patient 23

Amino-acid Matrix with NS mutations

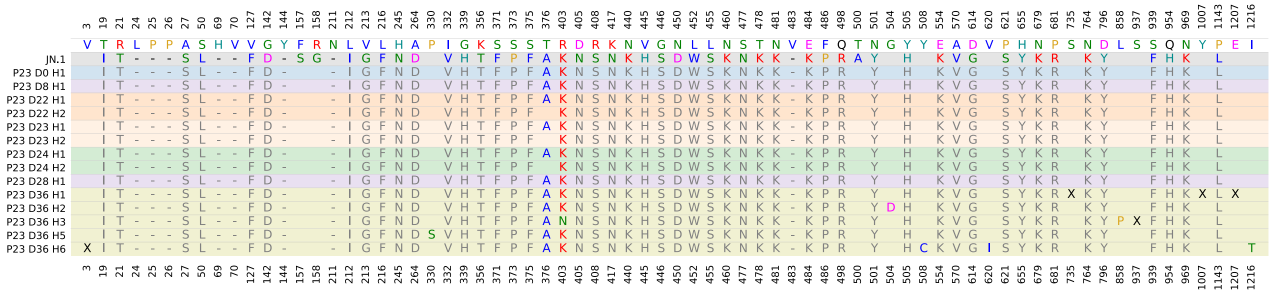


Divergence
Phylogeny Tree

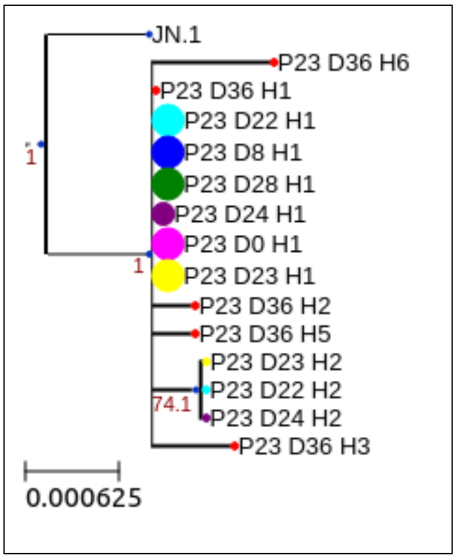


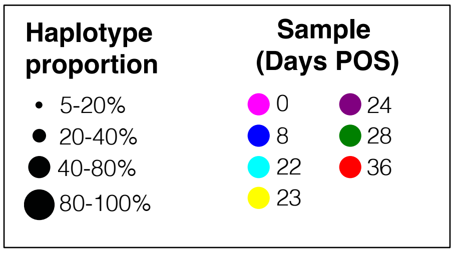


Molecular
Clock Tree

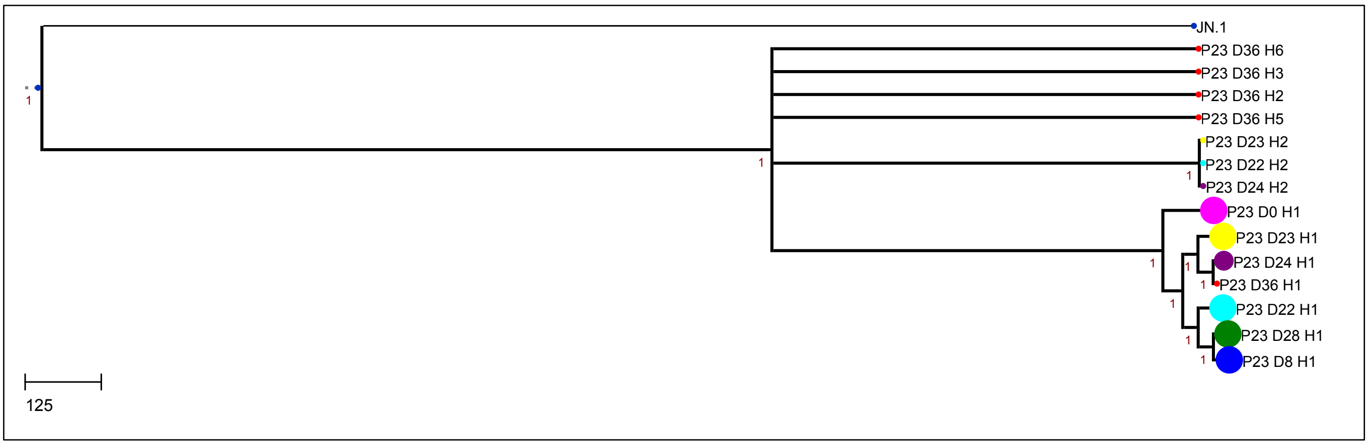

Supplemental Table 2

| | | Molecular Clock (s/s/y) | $R^2$ | (Residual mean)$^2$ | Fold Change related to contemporaneous VOC |
|---|---|---|---|---|---|
| Global whole genome estimates | | $5.8 \times 10^{-4}$ to $9.9 \times 10^{-4}$ (†) | | | |
| Lineage & Spike specific | | | | | |
| B.1 spike | | $5.08 \times 10^{-4}$ | 0.19 | $5.60 \times 10^{-8}$ | |
| B.1.1.7 spike | | $5.71 \times 10^{-4}$ | 0.13 | $1.29 \times 10^{-7}$ | |
| BA.1 spike | | $1.53 \times 10^{-3}$ | 0.22 | $3.76 \times 10^{-4}$ | |
| Anonymised ID / Lineage | | | | | |
| P1 | B.1.2 | $2.33 \times 10^{-3}$ | 0.47 | $3.34 \times 10^{-8}$ | |
| P2 | B.1 | $4.13 \times 10^{-3}$ | 0.87 | $4.48 \times 10^{-7}$ | 8.13 x |
| P3 | B.1.258 | $1.50 \times 10^{-3}$ | 0.54 | $8.93 \times 10^{-8}$ | |
| P4 | B.1.2 | $6.40 \times 10^{-3}$ | 0.29 | $1.13 \times 10^{-7}$ | |
| | B.1.177.18 | \\$3.00 \times 10^{-3}$ | 0.81 | $1.41 \times 10^{-8}$ | 2.95x |
| P5 | B.1.1.7 | $1.16 \times 10^{-3}$ | 0.20 | $6.92 \times 10^{-8}$ | |
| P6 | B.1.1.7 | $1.70 \times 10^{-3}$ | 0.20 | $2.60 \times 10^{-7}$ | |
| P7 | B.1.1.7 | $1.16 \times 10^{-3}$ | 0.42 | $6.88 \times 10^{-8}$ | |
| P8 | AY.4 | $1.32 \times 10^{-3}$ | 0.25 | $4.05 \times 10^{-7}$ | |
| P9 | BA.1.1 | $2.70 \times 10^{-3}$ | 0.26 | $1.22 \times 10^{-7}$ | |
| P10 | BA.2 | $1.15 \times 10^{-3}$ | 0.28 | $4.22 \times 10^{-8}$ | |
| P11 | BA.1.1 | $3.54 \times 10^{-3}$ | 0.92 | $8.87 \times 10^{-8}$ | 2,31x |
| P12 | BA.2 | $4.49 \times 10^{-4}$ | 0.13 | $5.70 \times 10^{-8}$ | |
| P13 | BA.2 | ND | NA | NA | |
| P14 | BA.2 | $1.88 \times 10^{-4}$ | 0.11 | $2.31 \times 10^{-8}$ | |
| P15 | BA.2 | $1.00 \times 10^{-3}$ | 1.0 | $5.43 \times 10^{-11}$ | 0.65x |
| P16 | BA.2 | $5.38 \times 10^{-4}$ | 0.17 | $4.36 \times 10^{-8}$ | |
| P17 | BA.2 | $1.20 \times 10^{-3}$ | 0.40 | $5.57 \times 10^{-8}$ | |
| P18 | BA.5.2 | $1.99 \times 10^{-4}$ | 0.19 | $4.28 \times 10^{-8}$ | |
| P19 | BA.5.2 | $2.05 \times 10^{-3}$ | 0.35 | $3.02 \times 10^{-6}$ | |
| P20 | BA.5.2 | $7.47 \times 10^{-3}$ | 0.34 | $2.24 \times 10^{-7}$ | |
| P21 | BQ.1.1 | $1.50 \times 10^{-3}$ | 0.40 | $1.88 \times 10^{-8}$ | |
| P22 | GE.1 | $1.26 \times 10^{-3}$ | 0.19 | $3.01 \times 10^{-7}$ | |
| P23 | JN.1 | $5.91 \times 10^{-3}$ | 0.76 | $5.30 \times 10^{-8}$ | 3.86x |

(†) Chaguza et al. 2023; Duchene et al. 2020

### Supplementary Figure 3

Detailed analyses of RBD changes and antibody escape

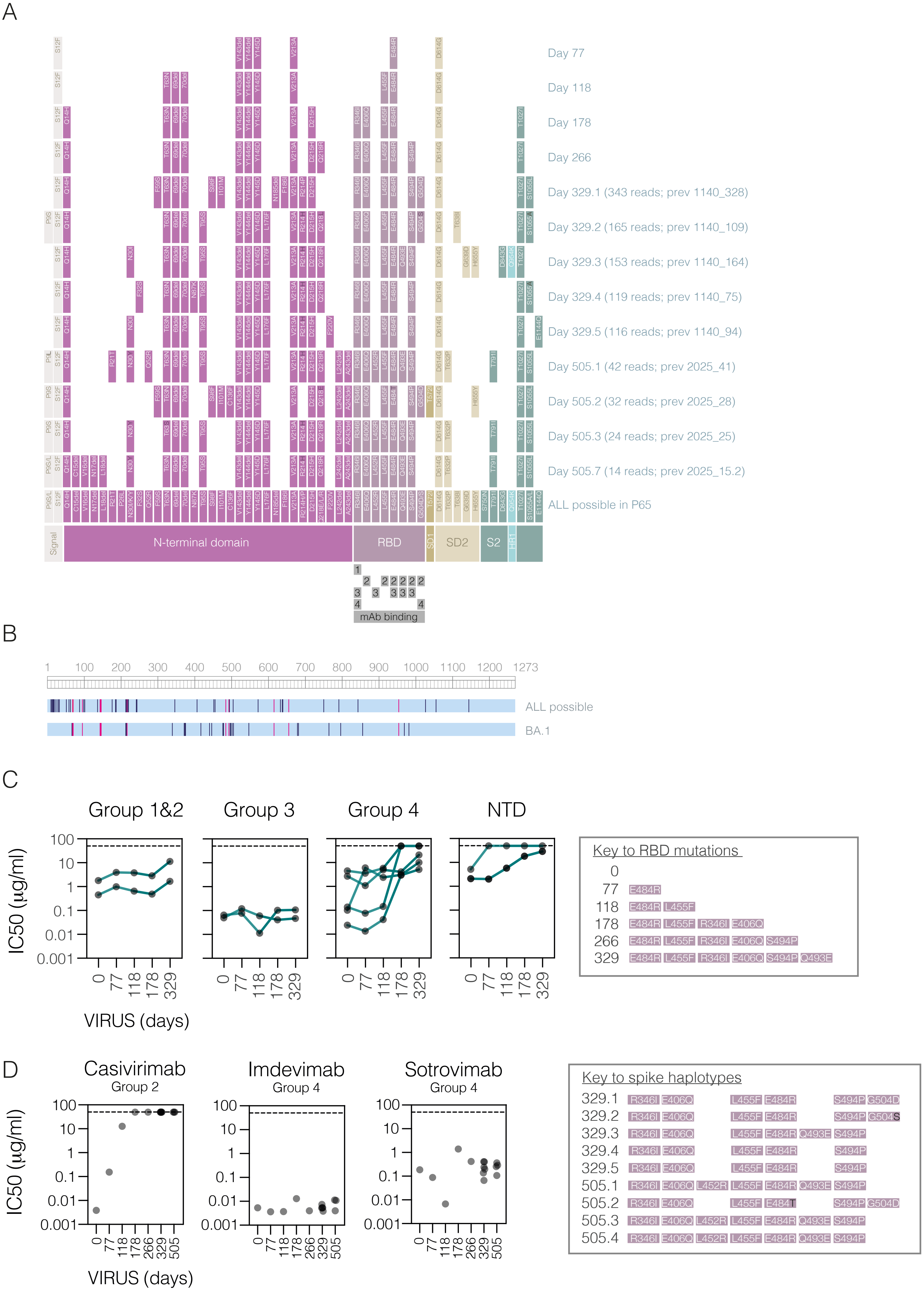
