## Supplemental Methods for "Antibody escape drives emergence of diverse spike haplotypes resembling variants of concern in persistent SARS-CoV-2 infections"

### Supplemental Methods
### Figure 1

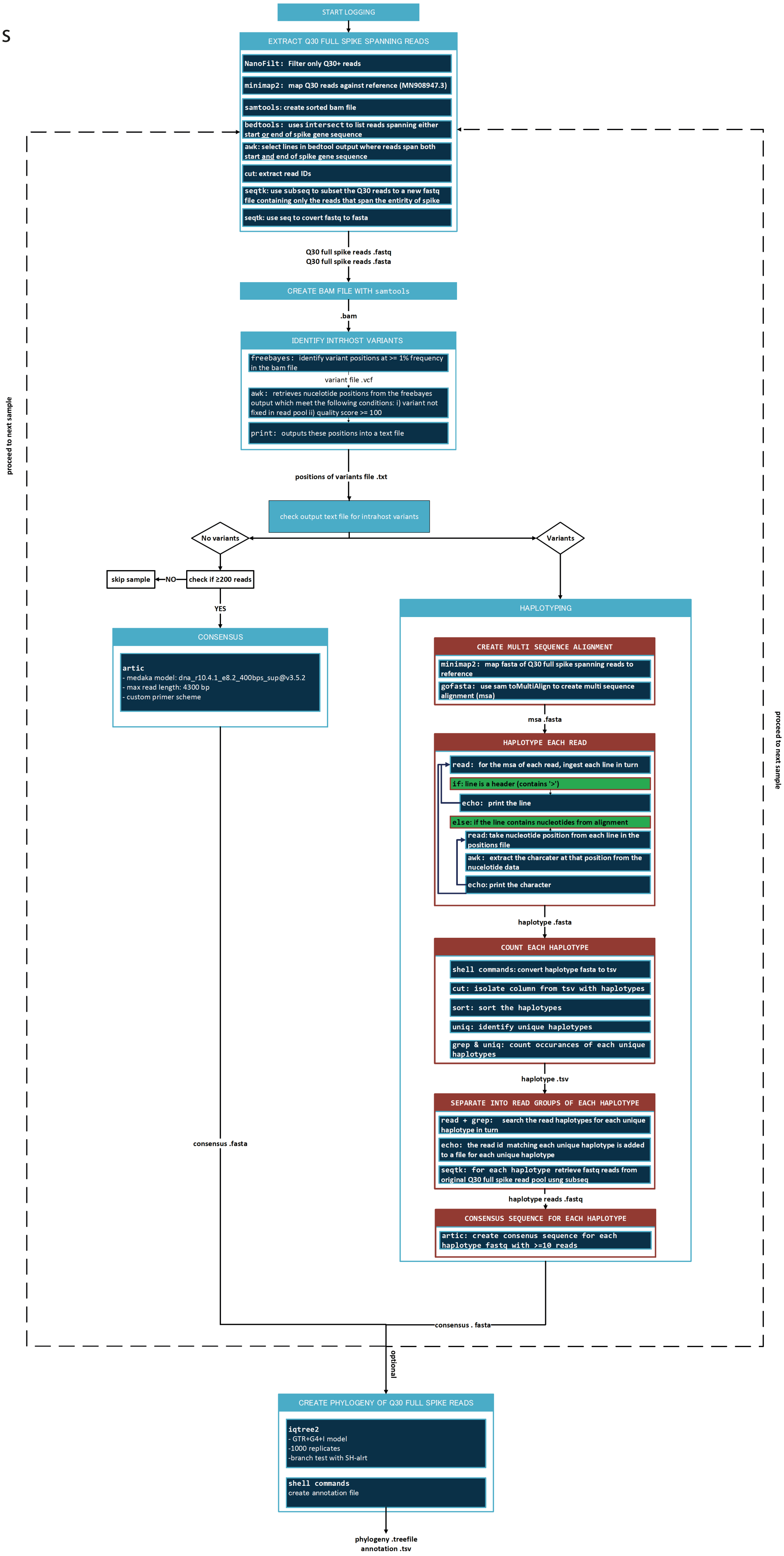

### Supplemental Methods
### Figure 2

## A

| Empirical mix; 100,000 viral RNA copies | | Mix identified by sequencing | | | | | |
|---|---|---|---|---|---|---|---|
| | | Replicate 1 | | Replicate 2 | | Replicate 3 | |
| **Alpha** | **Beta** | **Alpha** | **Beta** | **Alpha** | **Beta** | **Alpha** | **Beta** |
| **80%** | **20%** | 80.3% | 19.7% | 75.3% | 24.6% | 76.9% | 23.1% |
| **90%** | **10%** | 88.6% | 11.4% | 89.7% | 10.3% | 87.3% | 12.7% |
| **95%** | **5%** | 95.0% | 5.0% | 91.8% | 8.2% | 95.1% | 4.9% |
| **98%** | **2%** | 97.4% | 2.6% | 97.5% | 2.5% | 97.4% | 2.6% |
| **99%** | **1%** | 99.0% | 1.0% | 99.2% | 0.8% | 99.1% | 0.9% |
| **99.5%** | **0.5%** | 99.3% | 0.7% | 99.3% | 0.7% | 99.2% | 0.8% |
| **100%** | **0%** | not done | not done | 100% | 0% | 99.5% | 0.5% |

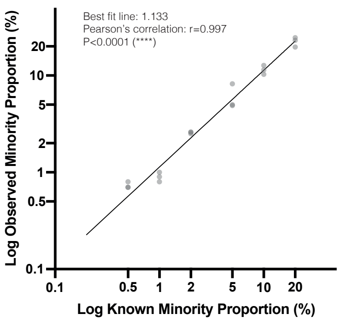


## B

| Viral Genome RNA copies | Q30 positive full-length spike reads | | | |
|---|---|---|---|---|
| | Replicate 1 | Replicate 2 | Replicate 3 | Replicate 4 |
| **1000** | 404 | 4339 | 4574 | 293 |
| **500** | 3333 | 1573 | 4997 | 1362 |
| **250** | 1934 | 2462 | 6514 | 714 |
| **100** | 825 | 3635 | 0 | 0 |

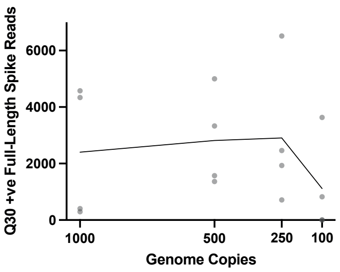


## C

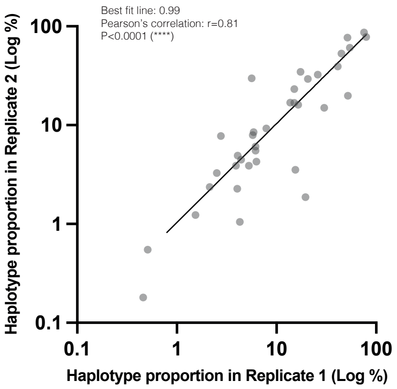

**Supplemental Methods Table 1** – Impact of down-sampling on the reproducibility of spike haplotype determination in clinical samples.

| SAMPLE | Q30+ full-spike reads | Haplotype 1 | | Haplotype 2 | | Haplotype 3 | | Haplotype 4 | | Haplotype 5 | | Haplotype 6 | | Haplotype 7 | |
|---|---|---|---|---|---|---|---|---|---|---|---|---|---|---|---|
| | | Reads | Percent (%) | Reads | Percent (%) | Reads | Percent (%) | Reads | Percent (%) | Reads | Percent (%) | Reads | Percent (%) | Reads | Percent (%) |
| **1** | 11336 | 8236 | 72.65 | 2908 | 25.65 | 83 | 0.73 | 49 | 0.43 | | | | | | |
| | 5000 | 3657 | 73.14 | 1273 | 25.46 | 36 | 0.72 | 18 | 0.36 | | | | | | |
| | 2500 | 1853 | 74.12 | 642 | 25.68 | 0 | 0.00 | 0 | 0.00 | | | | | | |
| | 1000 | 741 | 74.10 | 257 | 25.70 | 0 | 0.00 | 0 | 0.00 | | | | | | |
| | 500 | 371 | 74.20 | 128 | 25.60 | 0 | 0.00 | 0 | 0.00 | | | | | | |
| | 200 | 147 | 73.50 | 52 | 26.00 | 0 | 0.00 | 0 | 0.00 | | | | | | |
| **2** | 5074 | 3987 | 78.58 | 1067 | 21.03 | | | | | | | | | | |
| | 2500 | 1972 | 78.88 | 520 | 20.80 | | | | | | | | | | |
| | 1000 | 767 | 76.70 | 231 | 23.10 | | | | | | | | | | |
| | 500 | 384 | 76.80 | 113 | 22.60 | | | | | | | | | | |
| | 200 | 158 | 79.00 | 50 | 25.00 | | | | | | | | | | |
| **3** | 3892 | 3085 | 79.27 | 586 | 15.06 | 98 | 2.52 | 18 | 0.46 | 15 | 0.39 | 10 | 0.26 | 10 | 0.26 |
| | 2500 | 1965 | 78.60 | 385 | 15.40 | 70 | 2.80 | 11 | 0.44 | 0 | 0.00 | 0 | 0.00 | 0 | 0.00 |
| | 1000 | 773 | 77.30 | 166 | 16.60 | 27 | 2.70 | 0 | 0.00 | 0 | 0.00 | 0 | 0.00 | 0 | 0.00 |
| | 500 | 405 | 81.00 | 90 | 18.00 | 0 | 0.00 | 0 | 0.00 | 0 | 0.00 | 0 | 0.00 | 0 | 0.00 |
| | 200 | 156 | 78.00 | 41 | 20.50 | 0 | 0.00 | 0 | 0.00 | 0 | 0.00 | 0 | 0.00 | 0 | 0.00 |
| **4** | 2074 | 593 | 28.59 | 538 | 25.94 | 301 | 0.145 | 298 | 0.144 | 78 | 3.76 | 73 | 3.52 | 11 | 0.53 |
| | 1000 | 320 | 32.00 | 268 | 26.80 | 162 | 0.162 | 152 | 0.152 | 39 | 3.90 | 26 | 2.60 | 0 | 0.00 |
| | 500 | 151 | 30.20 | 131 | 26.20 | 83 | 0.166 | 81 | 0.162 | 21 | 4.20 | 20 | 4.00 | 0 | 0.00 |
| | 200 | 70 | 35.00 | 52 | 26.00 | 34 | 0.17 | 29 | 0.145 | 0 | 0.00 | 0 | 0.00 | 0 | 0.00 |
| **5** | 3465 | 1763 | 50.88 | 661 | 19.08 | 591 | 17.06 | 191 | 5.51 | 94 | 2.71 | 45 | 1.30 | | |
| | 2500 | 1305 | 52.20 | 482 | 19.28 | 437 | 17.48 | 143 | 5.72 | 70 | 2.80 | 32 | 1.28 | | |
| | 100 | 521 | 521.00 | 190 | 190.00 | 181 | 181.00 | 60 | 60.00 | 23 | 23.00 | 14 | 14.00 | | |
| | 500 | 264 | 52.80 | 97 | 19.40 | 79 | 15.80 | 31 | 6.20 | 17 | 3.40 | 0 | 0.00 | | |
| | 200 | 101 | 50.50 | 46 | 23.00 | 45 | 22.50 | 0 | 0.00 | 0 | 0.00 | 0 | 0.00 | | |

Supplemental Methods Table 2 – Replicate identification and quantification of full-spike haplotypes in clinical samples (n=9)

| SAMPLE | REPLICATE 1 | | | | REPLICATE 2 | | | |
|---|---|---|---|---|---|---|---|---|
| | Q30 Reads | Haplotype Rank | Reads | Haplotype Proportion | Q30 Reads | Haplotype Rank | Reads | Haplotype Proportion |
| **1** | 844 | 1 | 459 | 54.38 | 2030 | 1 | 1233 | 60.74 |
| **1** | 844 | 2 | 52 | 6.16 | 2030 | 3 | 112 | 5.52 |
| **1** | 844 | 3 | 52 | 6.16 | 2030 | 2 | 123 | 6.06 |
| **1** | 844 | 4 | 34 | 4.03 | 2030 | 6 | 46 | 2.27 |
| **1** | 844 | 5 | 33 | 3.91 | 2030 | 4 | 79 | 3.89 |
| **1** | 844 | 6 | 18 | 2.13 | 2030 | 5 | 48 | 2.36 |
| **1** | 844 | 7 | 13 | 1.54 | 2030 | 7 | 25 | 1.23 |
| **1** | 844 | 8 | 11 | 1.30 | 2030 | 8 | 0 | 0.00 |
| **2** | 1972 | 1 | 512 | 25.96 | 1827 | 1 | 591 | 32.35 |
| **2** | 1972 | 2 | 406 | 20.59 | 1827 | 2 | 536 | 29.34 |
| **2** | 1972 | 3 | 325 | 16.48 | 1827 | 4 | 294 | 16.09 |
| **2** | 1972 | 4 | 271 | 13.74 | 1827 | 3 | 309 | 16.91 |
| **2** | 1972 | 5 | 124 | 6.29 | 1827 | 5 | 78 | 4.27 |
| **2** | 1972 | 6 | 104 | 5.27 | 1827 | 6 | 71 | 3.89 |
| **2** | 1972 | 7 | 10 | 0.51 | 1827 | 8 | 10 | 0.55 |
| **2** | 1972 | no | 0 | 0.00 | 1827 | 7 | 12 | 0.66 |
| **3** | 388 | 1 | 200 | 51.55 | 480 | 1 | 369 | 76.88 |
| **3** | 388 | 2 | 117 | 30.15 | 480 | 2 | 72 | 15.00 |
| **3** | 388 | 3 | 60 | 15.46 | 480 | 3 | 17 | 3.54 |
| **4** | 3892 | 1 | 3085 | 79.27 | 8215 | 1 | 6419 | 78.14 |
| **4** | 3892 | 2 | 586 | 15.06 | 8215 | 2 | 1383 | 16.84 |
| **4** | 3892 | 3 | 98 | 2.52 | 8215 | 3 | 269 | 3.27 |
| **4** | 3892 | 4 | 18 | 0.46 | 8215 | 5 | 15 | 0.18 |
| **5** | 11336 | 1 | 3158 | 27.86 | 5185 | 1 | 8236 | 158.84 |
| **5** | 11336 | 2 | 1804 | 15.91 | 5185 | 2 | 2908 | 56.08 |
| **5** | 11336 | 3 | 183 | 1.61 | 5185 | 3 | 0 | 0.00 |
| **5** | 11336 | 4 | 21 | 0.19 | 5185 | 4 | 0 | 0.00 |
| **6** | 3393 | 1 | 1763 | 51.96 | 1710 | 3 | 340 | 19.88 |
| **6** | 3393 | 2 | 661 | 19.48 | 1710 | 6 | 32 | 1.87 |
| **6** | 3393 | 3 | 591 | 17.42 | 1710 | 1 | 591 | 34.56 |
| **6** | 3393 | 4 | 191 | 5.63 | 1710 | 2 | 510 | 29.82 |
| **6** | 3393 | 5 | 94 | 2.77 | 1710 | 5 | 133 | 7.78 |
| **6** | 3393 | 6 | 0 | 0.00 | 1710 | 4 | 16 | 0.94 |
| **7** | 747 | 1 | 335 | 44.85 | 2189 | 1 | 1160 | 52.99 |
| **7** | 747 | 2 | 59 | 7.90 | 2189 | 2 | 202 | 9.23 |
| **7** | 747 | 3 | 44 | 5.89 | 2189 | 3 | 186 | 8.50 |
| **7** | 747 | 4 | 43 | 5.76 | 2189 | 4 | 173 | 7.90 |
| **7** | 747 | 5 | 33 | 4.42 | 2189 | 5 | 98 | 4.48 |
| **7** | 747 | 6 | 32 | 4.28 | 2189 | 9 | 23 | 1.05 |
| **8** | 5074 | 1 | 3828 | 75.44 | 4605 | 1 | 3987 | 86.58 |
| **8** | 5074 | 2 | 766 | 15.10 | 4605 | 2 | 1067 | 23.17 |
| **9** | 909 | 1 | 486 | 53.47 | 1259 | 1 | 1658 | 131.69 |
| **9** | 909 | 2 | 375 | 41.25 | 1259 | 2 | 494 | 39.24 |
| **9** | 909 | 3 | 37 | 4.07 | 1259 | 3 | 62 | 4.92 |
